# Arenavirus nucleoprotein localizes to mitochondria

**DOI:** 10.1101/2020.11.06.370825

**Authors:** Francesca Baggio, Udo Hetzel, Lisbeth Nufer, Anja Kipar, Jussi Hepojoki

## Abstract

Viruses need cells to replicate and, therefore, ways to counteract the host’s immune response. Mitochondria play central roles in mediating innate immunity, hence some viruses have developed mechanisms to alter mitochondrial functions. Herein we show that arenavirus nucleoprotein (NP) enters the mitochondria of infected cells and affects their morphological integrity. We initially demonstrate electron-dense inclusions within mitochondria of reptarenavirus infected cells and hypothesized that these represent viral NP. Software predictions then serve to identify a putative N-terminal mitochondrial targeting signal (MTS) in arenavirus NPs; however, comparisons of wild-type and N-terminus mutated NPs suggest MTS-independent mitochondrial entry. NP does not enter isolated mitochondria, indicating that translocation requires additional cellular factors or conditions. Immune electron microscopy finally confirms the presence of NP within the mitochondria both *in vitro* and in infected animals. We hypothesize that mitochondria targeting might complement the known interferon antagonist functions of NP or alter the cell’s metabolic state.

## INTRODUCTION

Viruses are obligate cell parasites that have adapted specific strategies to either promote or inhibit various host cell functions for their own benefit. To hijack the host cell, viruses can target specific subcellular structures and alter their morphology, or can affect host protein levels and/or their trafficking^1^. To counteract the host defense, viruses can interfere with the functions of cellular compartments that contribute to antiviral signalling, such as mitochondria, peroxisomes, endoplasmic reticulum (ER), lipid droplets and the nucleus^1,2^.

Mitochondria, in addition to their essential role in metabolism, regulate apoptosis^3^ and control both immune and inflammatory responses^4^. Viruses can perturb mitochondria in several ways, e.g. by altering metabolic pathways, by inducing fusion or fission to promote mitochondrial autophagy (mitophagy) or affect mitochondrial distribution within the cell^5^. Mitochondria play an essential role in the cell’s antiviral response. Their outer membrane houses the mitochondrial antiviral-signaling protein (MAVS) which mediates both the type I interferon (IFN) response and pro-inflammatory pathways^6,7^. Retinoic acid-inducible gene I (RIG-I) and melanoma differentiation-associated protein 5 (MDA-5) are pattern recognition receptors (PRRs) that sense dsRNAs in the cytosol and, following ligand binding, induce MAVS aggregation^8^. This in turn leads to nuclear translocation of IFN regulatory factor 3 (IRF-3) and subsequent activation of type I IFN signaling pathways, which contribute to the cell-mediated immunity^8^. In addition, MAVS participates in the regulation of apoptosis^9,10^. The dynamic mitochondrial fusion and fission cycle, in combination with mitophagy, promotes or hampers the MAVS-mediated innate immunity^11^. Mitochondrial fusion facilitates downstream signaling cascades of MAVS and its interaction with the ER-associated innate immunity molecule stimulator of interferon genes (STING), whereas mitochondrial fission has opposite effects^12^. Mitochondria are also involved in adaptive immunity via generation of reactive oxygen species (ROS), contributing to T cell activation, and via mitochondrial metabolism, regulating CD4+/CD8+ (memory *vs* effector) T cell differentiation^4^. Thus, targeting mitochondria or signal molecules up- or downstream of MAVS is a general immune evasion strategy of viruses^13^.

The family *Arenaviridae* comprises four genera, *Mammarenavirus*, *Reptarenavirus*, *Hartmanivirus* and *Antennavirus*^14^. Arenaviruses have a single-stranded bisegmented RNA genome, except for antennaviruses that have a trisegmented genome^15^. The large (L) segment of mamm- and reptarenaviruses encodes the Z protein (ZP) and the RNA-dependent RNA polymerase (RdRp), and the small (S) segment encodes the glycoprotein precursor (GPC) and the nucleoprotein (NP)^15^. Hartmaniviruses and, likely, antennaviruses, lack the ZP^16^. The mammarenavirus NP and ZP both suppress the host’s innate immunity by antagonizing the type I IFN response^17^. The C-terminal domain of NP harbors an exoribonuclease activity that suppresses the type I IFN response by degrading dsRNA replication products^18^ and blocking the dsRNA-activated protein kinase (PKR) signaling pathway^19^. The NP directly binds to RIG-I, MDA5^20^, and inhibitor of nuclear factor kappa-B kinase subunit epsilon (IKK∊), a factor acting downstream of MAVS^21^, thus inhibiting the nuclear translocation and transcriptional activity of IRF-3^22,23^ and nuclear factor kappa-B (NF-κB)^24^. The ZPs of some mammarenaviruses also prevent MAVS activation by interacting with RIG-I or MDA-5^25,26^. Furthermore, the ZP of lymphocytic choriomeningitis virus (LCMV) causes re-localization of promyelocytic leukemia nuclear bodies, IFN-induced nuclear components, to the cytoplasm, thus counteracting the host’s antiviral response^1,27,28^.

Reptarenaviruses were discovered in the early 2010s in captive constrictor snakes with boid inclusion body disease (BIBD)^29–31^. BIBD manifests by the formation of electron-dense cytoplasmic inclusion bodies (IBs) in most cell types of affected snakes^32,33^. The IBs, comprising mainly reptarenavirus NP^29,31,34^, appear not to be cytopathic^31,35^. However, snakes with BIBD often die from secondary bacterial, fungal or protozoal infections, or neoplastic processes^36,37^, suggesting that reptarenaviruses cause immunosuppression^38,39^. This is supported by the fact that snakes with BIBD carry IBs in leukocytes and myelopoietic cells and often exhibit a weak antibody response against reptarenavirus NP^31,38,40^.

Herein, by combining ultrastructural imaging and molecular biology techniques we demonstrate that reptarenavirus NP induces IB formation within the mitochondria of host cells. We have identified a putative mitochondrial translocation signal (MTS) in rept- and mammarena-but not in hartmanivirus NPs. Using recombinant NPs of these arenaviruses, we show that mitochondrial translocation is a common feature of arenavirus NPs. However, different from most proteins that translocate into mitochondria, the putative MTSs of the studied NPs remain uncleaved in reptilian and mammalian cells. Moreover, while mutations to the putative MTSs of rept- and mammarenavirus NPs appear to reduce their oligomerization potential, they do not affect mitochondrial translocation. These results indicate that putative arenavirus MTSs might contribute to IB formation but that the mitochondrial translocation might be mediated *via* an alternative, not yet identified NP sequence or pathway. In addition, arenaviral NPs are not imported into isolated mitochondria, suggesting the necessity of additional cell components or conditions that are not preserved in this *in vitro* system. Finally, utilizing immune EM on the brain of boas euthanized due to BIBD, we demonstrate *in situ* that mitochondrial inclusions comprising NP occur also *in vivo*.

## RESULTS

### Reptarenavirus NP forms inclusions within mitochondria

In the attempt to identify any subtle cytopathic effects of reptarenaviruses, we investigated the boid kidney (I/1 Ki) cell line infected with University of Giessen virus 1 (UGV-1) by transmission electron microscopy (TEM). At three days post-infection (dpi) reptarenavirus infected cells demonstrated the characteristic electron-dense round to oval IBs with smooth boundaries^31^, but also less electron-dense IBs with rough edges (Fig. 1a-c). The mitochondria exhibited pronounced vacuolization and disruption of the matrix with loss of cristae structures and occasional evidence of intramitochondrial IB formation (Fig. 1a-c).

**Figure 1.**
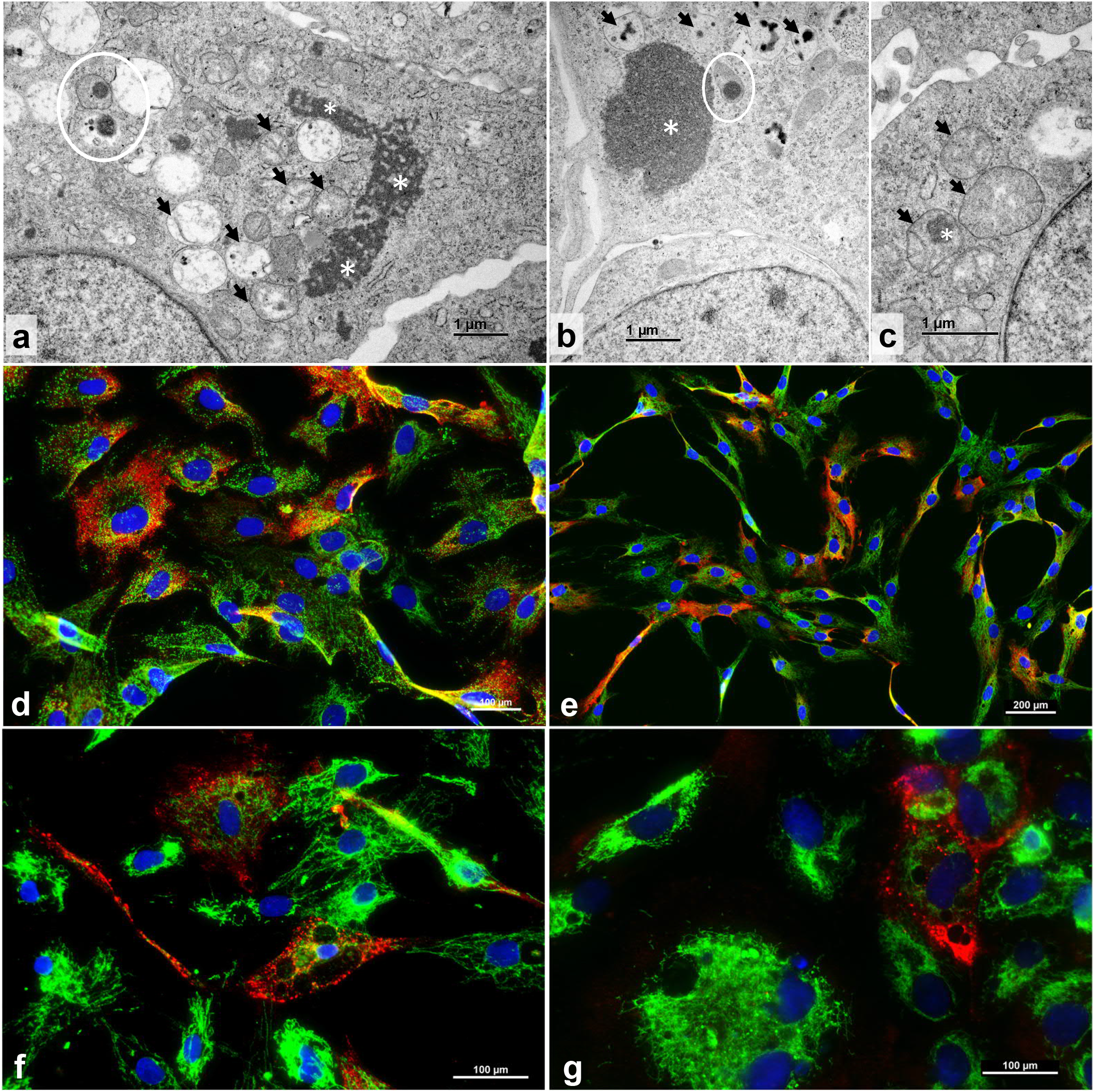
Transmission electron microscopy (TEM) and immunofluorescence (IF) analyses of reptarenavirus infected boid cell lines. **(a-b)** TEM, permanent cell culture derived from *B. constrictor* kidney (I/1 Ki), infected with UGV-1, at three dpi. **(a)** Large irregular cytoplasmic inclusion body (IB; asterisks), vacuolated mitochondria (arrows) and partly ruptured mitochondrion with electron-dense IBs in the matrix (circle). **(b)** Large electron-dense cytoplasmic IB (asterisk), one mitochondrion with IB in the matrix (circle), and several vacuolar structures consistent with vacuolated mitochondria with small IBs (arrows). **(c)** Swollen mitochondria with finely granular disintegrated matrix (arrows) and one IB (asterisk). **(d-g)** IF of permanent tissue culture lines derived from *B. constrictor* brain (V/4 Br) **(d)**, kidney (I/1 Ki) **(e)**, lungs (V/4 Lu) **(f)** and liver (V/1 Liv) **(h)**, infected with the “equimolar” mix of UGV-1, UHV-1 and ABV-1, at three dpi, showing some infected cells with a less pronounced mitochondrial staining. Reptarenavirus nucleoprotein (NP) in red (AlexaFluor 594 goat anti-rabbit), mitochondrial marker (mtCO2) in green (AlexaFluor 488 goat anti-mouse), nuclei in blue (DAPI). See also Supplementary Figure 1.

Since IBs are mainly composed of reptarenavirus NP^29,31,34^, we further investigated the relation between reptarenaviral NP and the mitochondrial network and performed immunofluorescence (IF) analyses on four different *Boa constrictor* cell lines, derived from brain (V/4 Br), kidney (I/1 Ki), lung (V/4 Lu), and liver (V/1 Liv). To mimic the naturally occurring reptarenavirus co-infections^41–43^, the cell lines were inoculated with an “equimolar” mix (as determined by qRT-PCR; Supplementary Table 1) of Aurora borealis virus 1 (ABV-1), University of Helsinki virus 1 (UHV-1), and UGV-1. Interestingly, at three dpi, while IF did not reveal a definite co-localization of NP and the mitochondrial marker (mtCO2), the NP staining pattern appeared to follow the organization of the mitochondrial network; also, in some infected cells, staining for mtCO2 appeared less pronounced, suggesting a reduction in mitochondria (Fig.1d-g, Supplementary Fig.1).

### *In silico* analysis of arenavirus NP’s cellular localization

Many newly synthesized proteins destined to the mitochondria carry a mitochondrial targeting signal (MTS), an N-terminal positively charged amphipathic helix of 15-70 amino acid residues^44^. We used two online prediction tools, Target P 1.1 (http://www.cbs.dtu.dk/services/TargetP-1.1/index.php) and Mitoprot II (https://ihg.gsf.de/ihg/mitoprot.html) to study arenavirus NPs. The analysis revealed a putative MTS in all tested reptarenavirus NPs, UGV-1, UHV-1, ABV-1, Golden Gate virus 1 (GGV-1), Tavallinen suomalainen mies virus 1 (TSMV-1), Rotterdam reptarenavirus (ROUT) and California reptarenavirus (CASV); with Target P1.1 the predictions ranged from 33.5 to 59.5%, and with Mitoprot II from 84.8 to 97.6% (Table 1). However, TargetP 1.1 predicted only UGV-1, TMSV-1, and ROUT NPs to have an MTS cleavage site between residues 7 and 8, and between 47 and 48 for ABV-1 NP. Mitoprot II predicted a cleavage site only for ROUT NP, between residues 29 and 30 (Table 1). We then studied whether the NPs of viruses from the other genera of the family *Arenaviridae* would produce similar predictions. Indeed, as summarized in Table 1, many mammarenavirus NPs contain a putative MTS, however, neither hartmani-nor antennavirus NPs contain an MTS by prediction (Table 1).

**Table 1.**
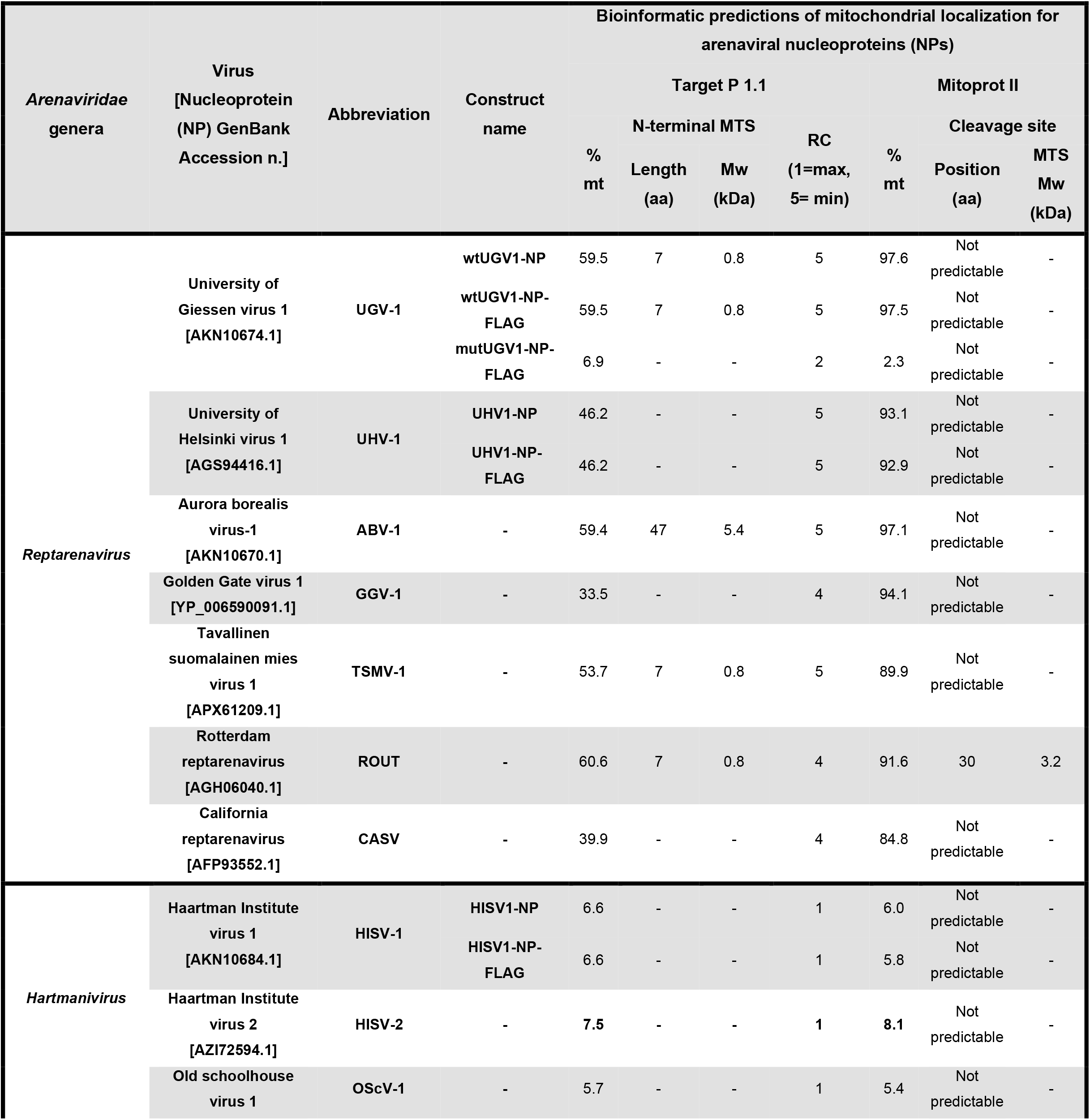
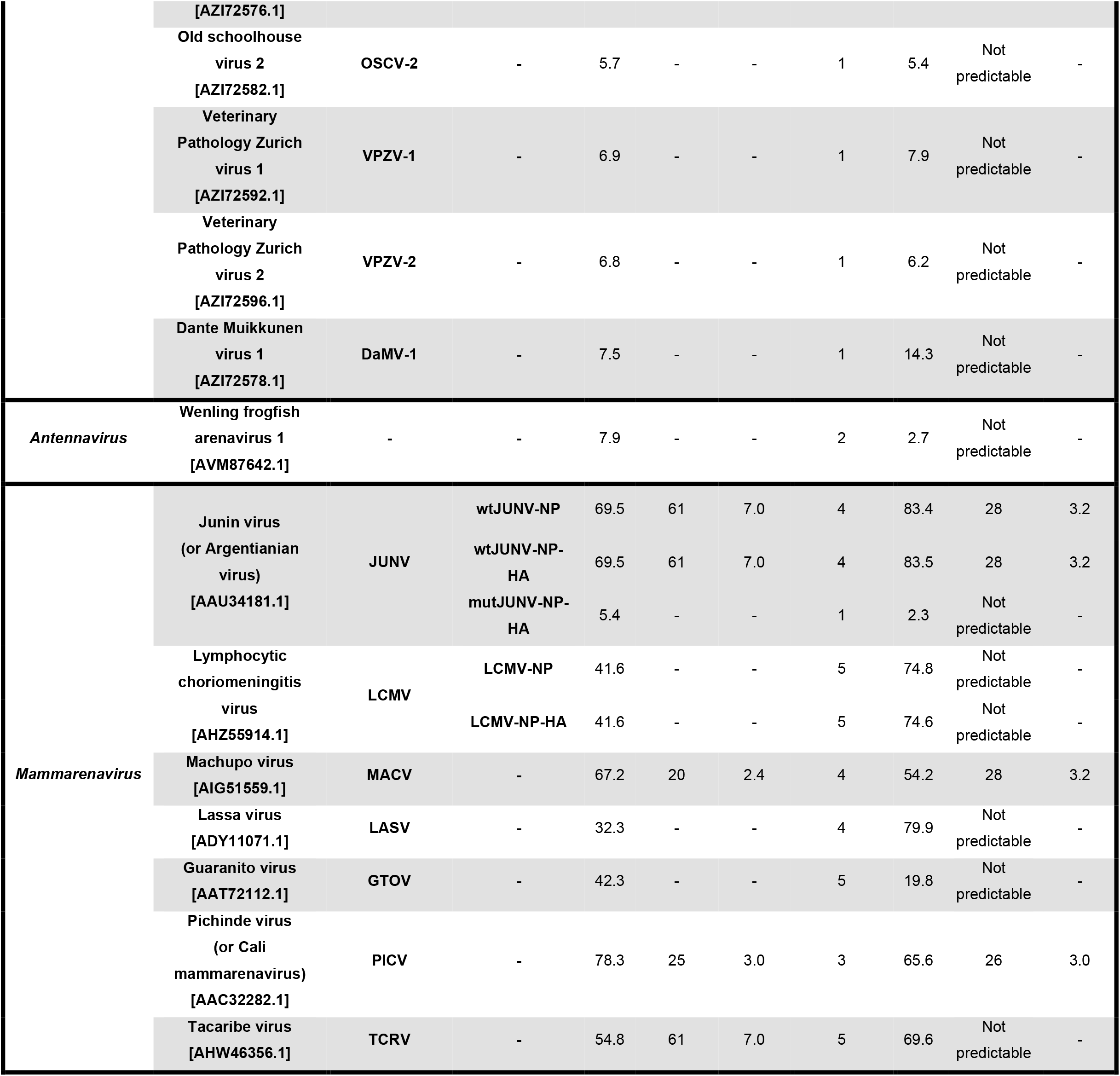
*In silico* predictions of mitochondrial localization for different arenaviral NPs.

### Mitochondrial translocation of NP is MTS-independent

Because the MTS is a positively charged amphipathic helix^44^, we generated recombinant expression constructs in which the positively charged interface is replaced by negative charges. We used Jpred, a protein secondary structure prediction tool^45^, to confirm that the introduced mutations do not alter the helical structure of the putative MTS. Target P 1.1 and Mitoprot II served to demonstrate a loss of mitochondrial translocation potential of the NPs with mutated MTS (Table 1). MTS mutations were introduced to the NPs of UGV-1 and JUNV, to generate FLAG- and HA-tagged constructs: wtUGV1-NP-FLAG, mutUGV1-NP-FLAG, wtJUNV-NP-HA, and mutJUNV-NP-HA (Fig. 2a,b). UHV-1, LCMV, and HISV-1 NP constructs, UHV1-NP-FLAG, LCMV-NP-HA, and HISV1-NP-FLAG, served as controls. At three days post-transfection (dpt) IF analysis of *Boa constrictor* V/4 Br cells transfected with wtUGV1-NP-FLAG, mutUGV1-NP-FLAG, wtJUNV-NP-HA, and mutJUNV-NP-HA constructs showed that the wtNPs yielded a punctate staining pattern commonly seen in reptarenavirus infected cells, whereas the MTS-mutated NPs induced a diffuse staining pattern (Fig. 2c-f). In addition, V/4 Br cells transfected with UHV1-NP-FLAG bearing the putative MTS (Table 1) consistently exhibited the punctate NP staining pattern (Fig. 2c,d). In contrast, LCMV NP, which also bears a putative MTS (Table 1), yielded a diffuse staining pattern (Fig. 2e,f). Interestingly, HISV-1 NP, devoid of the MTS-like sequence (Table 1), was either diffusely distributed in the cells or formed perinuclear tubular structures (Fig. 2c,d), confirming our earlier IF, TEM and immune EM findings in *Boa constrictor* I/1 Ki cells^16^. These findings suggest that the putative MTS of many arenavirus NPs plays a role in the oligomerization/aggregation and potentially contributes to IB formation of both rept- and mammarenaviruses. To further investigate the fate of the NP’s N-terminal region, we produced an antiserum against the putative MTSs of rept- and mammarenavirus NPs. IF staining using the anti-MTS antibody produced a diffuse staining pattern in cells transfected with the mutNP constructs, and, curiously, cells expressing wtNPs, including LCMV NP, showed no staining (Supplementary Fig. 2a-d). Because only mutNPs were detected with anti-MTS, we speculate that the oligomerization of wtNPs renders the putative MTS inaccessible and thus prevents IF staining using antibodies targeting this region (Supplementary Fig. 2a-d). Exceptionally, LCMV NP did neither produce larger cellular aggregates (Fig. 2e,f) nor induce staining with anti-MTS (Supplementary Fig. 2c,d), leading us to hypothesize that it is rather the tertiary or quaternary structure of NPs than aggregate formation *per se* that renders the N-terminus inaccessible to epitope recognition. When we included the mitochondrial marker mtCO2 to the IF analyses to locate the arenavirus NPs in relation to mitochondria, there was again no clear evidence of co-localization, neither for wt nor mutNP variants (Fig. 2c-f). However, the NP staining occasionally appeared to follow the mitochondrial network (Fig. 2c-f). These results are in line with the ultrastructural observations that not all mitochondria develop inclusions, and that mitochondria containing NP often lose their structural integrity.

**Figure 2.**
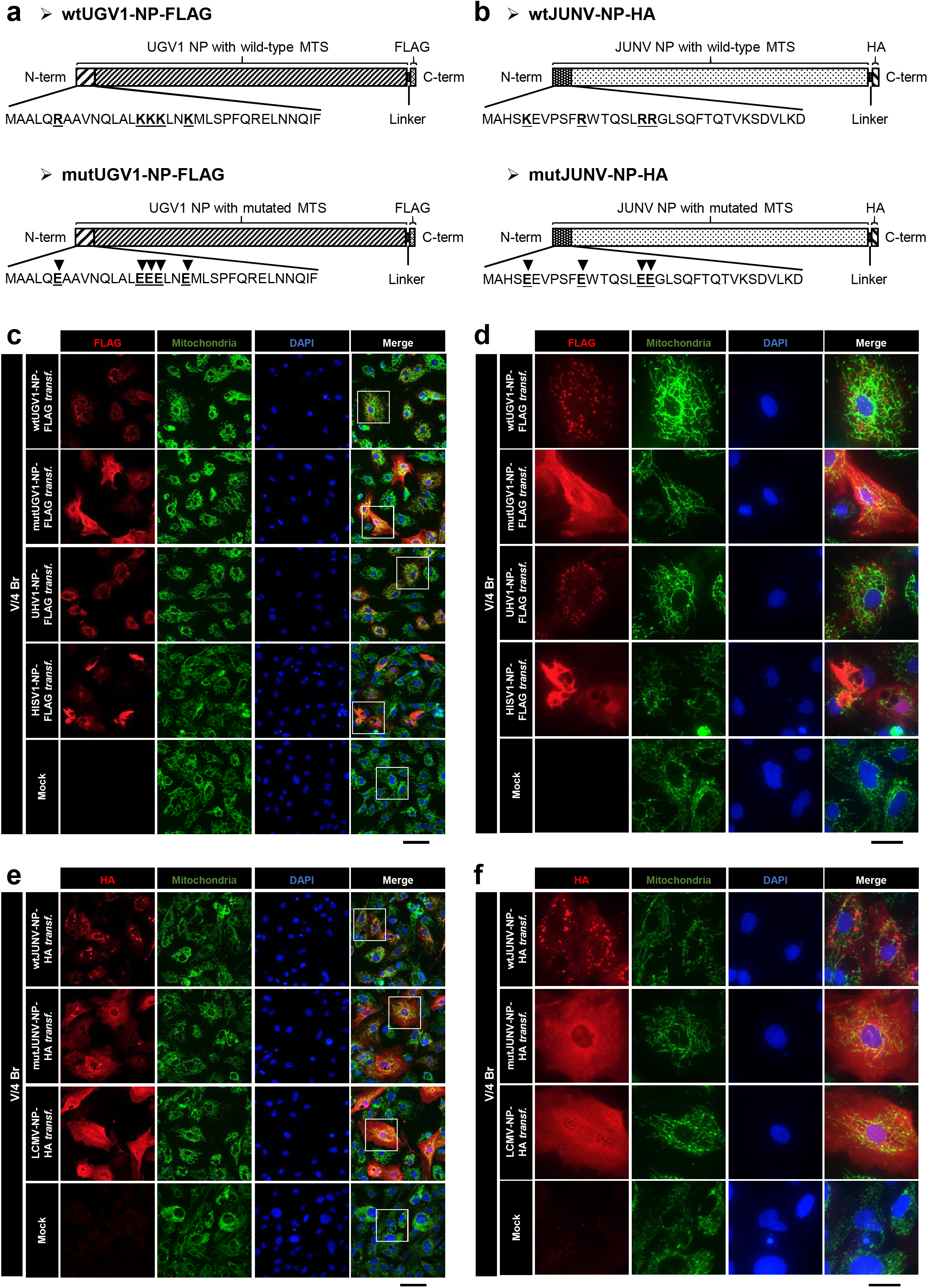
IF analyses of the expression pattern of different arenaviral NPs in transfected boid cells. **(a-b)** Schematic representation of the wild-type (wt) and the mutated (mut) versions of the of UGV-1 NP **(a)** and JUNV NP **(b)** expressed from a pCAGGS-FLAG or pCAGGS-HA construct, respectively, and used for transfections. Black arrowheads in the putative MTSs (first N-terminal 34 amino acids shown) indicate the positions of the amino acid substitutions of the mut versions compared to the corresponding wt sequences. Both wt and mut UGV-1 NPs are fused in frame with a C-terminal FLAG tag, separated by a linker sequence **(a)**. Both wt and mut JUNV NPs are fused in frame with a C-terminal HA tag, separated by a linker sequence **(b)**. **(c-f)** Double IF images of *Boa constrictor* V/4 Br cells transfected with a construct expressing either wt or mutUGV1-NP-FLAG, UHV1-NP-FLAG, or HISV1-NP-FLAG **(c,d)**, and wt or mutJUNV-NP-HA, or LCMV-NP-HA **(e,f)** at three dpt. Non-transfected (Mock) cells served as controls. **(c,d)** The panels from left: FLAG tag in red (AlexaFluor 594 goat anti-rabbit), mitochondrial marker (mtCO2) in green (AlexaFluor 488 goat anti-mouse), nuclei in blue (DAPI), and a merged image. **(e,f)** The panels from left: HA tag in red (AlexaFluor 594 goat anti-rabbit), mitochondrial marker (mtCO2) in green (AlexaFluor 488 goat anti-mouse), nuclei in blue (DAPI), and a merged image. White squares in the merged images in **(c,e)** define the magnified areas in **(d,f)**. Scale bars: 500 μm **(c,e)** and 200 μm **(d,f)**. See also Supplementary Figure 2.

Following mitochondrial translocation, specific peptidases cleave the MTS of proteins destined to the mitochondrial matrix^46^. To investigate whether the putative MTSs of arenavirus NPs are cleaved, we studied V/4 Br cells transfected with wtUGV1-NP-FLAG, mutUGV1-NP-FLAG, wtJUNV-NP-HA and mutJUNV-NP-HA as well as UHV1-NP-FLAG, LCMV-NP-HA and HISV1-NP-FLAG at three dpt by immunoblotting. Tubulin and mitofusin-2 (MFN2) served as cytosolic and mitochondrial markers, respectively (Ponceau S staining of the membranes, Supplementary Fig. 3a-d). Both wt and mutNPs produced a single main band with anti-FLAG or anti-HA antibodies in whole-cell lysates as well as in isolated mitochondria; based on the estimated molecular weight (approx. 63-68 kDa) this represents non-cleaved NP (Fig. 3a-d) which was also confirmed for HISV-1 NP by employing anti-Hartmani NP antiserum (Fig. 3d). To demonstrate that the N-terminus remains intact, we performed immunoblotting using the anti-MTS antiserum, thereby confirming that the NPs remain non-cleaved (Fig. 3a-c). Furthermore, the immunoblotting revealed that both non-cleaved wt and mutNPs migrated in the mitochondrial fractions (Fig. 3a-d). To rule out the possibility that the NPs are synthesized within the mitochondria, we performed an *in silico* analysis of the open reading frame (ORF) encoding the NPs using Translate (available at https://web.expasy.org/translate/). As the codon usage varies between both species and organelles, we compared the sizes of predicted translation products from the NP ORFs by the cytosolic ribosomes to those obtained from mitoribosomes. The analysis of UGV-1, UHV-1, HISV-1, JUNV, and LCMV NP translation produced the expected 62-67 kDa protein products only when the standard translation code was used (Supplementary Table 2). The vertebrate mitochondrial code interprets AGA and AGG as termination codons instead of Arginin^47^, therefore, the translation produced truncated peptides of only 0.5-4 kDa (Supplementary Table 2). The findings indicate that cytosolic ribosomes mediate NP translation, and that the mitochondrial translocation occurs post-translationally.

**Figure 3.**
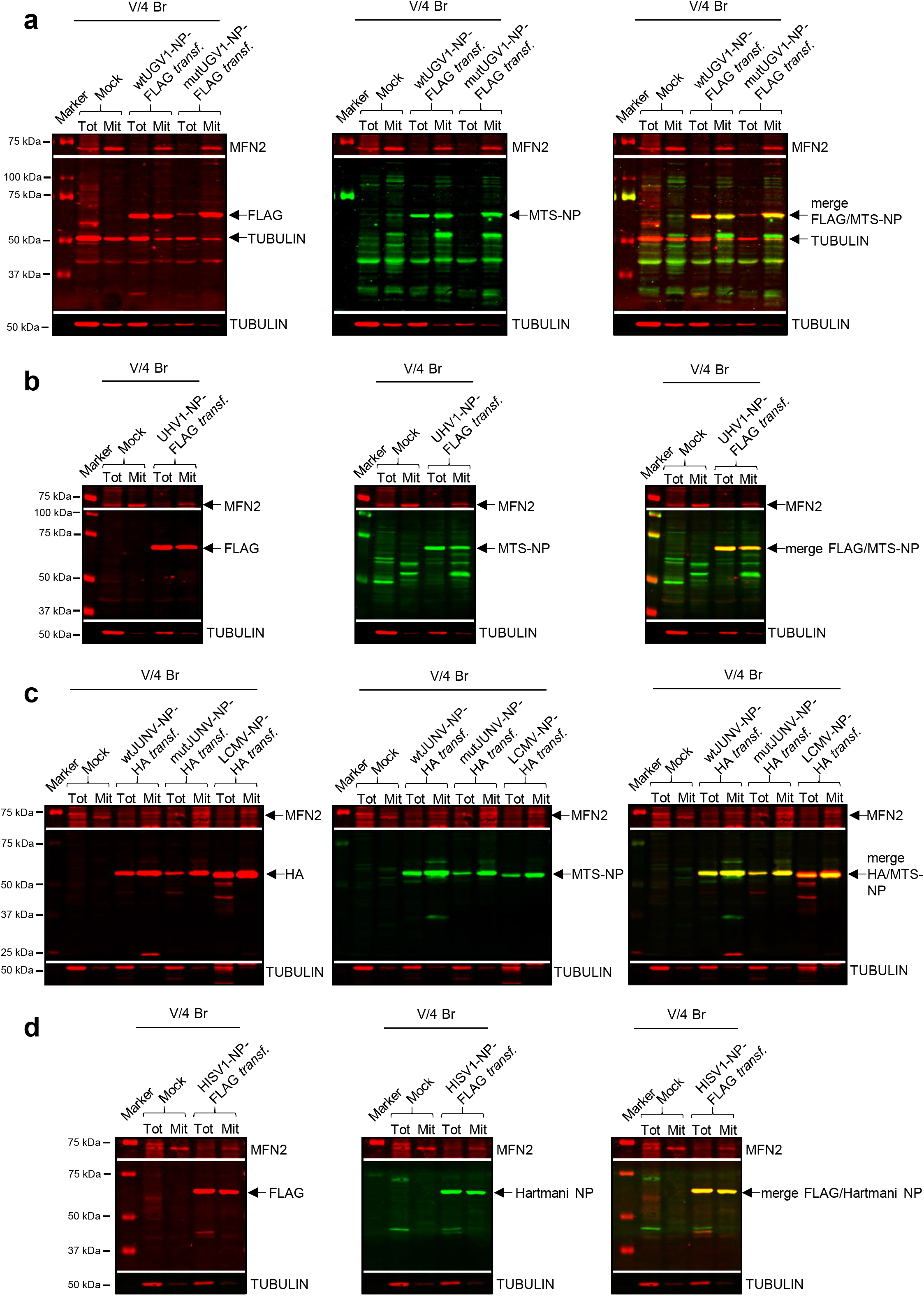
Immunoblotting studies on boid cells transfected with different arenaviral NPs. **(a-d)** Immunoblotting analyses of whole-cell lysates (Tot) and mitochondrial preparations (Mit) obtained from *Boa constrictor* V/4 Br cells transfected with constructs expressing wt or mutUGV1-NP-FLAG **(a)**, UHV1-NP-FLAG **(b)**, wt or mutJUNV-NP-HA, or LCMV-NP-HA **(c)** and HISV1-NP-FLAG **(d)**, all at three dpt. Non-transfected (Mock) samples were used as negative controls. 20 μg **(a,b)** or 12 μg **(c,d)** protein samples were loaded on standard SDS-PAGE gels followed by immunoblotting analyses. Tubulin and MFN2 (both in red) were used as cytosolic and mitochondrial marker, respectively. Arenavirus NPs (63-68 kDa) are indicated (black arrows). **(a,b)** Left panels: FLAG tag in red (IRDye 680RD Donkey anti-mouse); middle panels: MTS-NP in green (IRDye 800CW Donkey anti-rabbit); right panels: merged image. **(c)** Left panel: HA tag in red (IRDye 680RD Donkey anti-mouse); middle panel: MTS-NP in green (IRDye 800CW Donkey anti-rabbit); right panel: merged image. **(d)** Left panel: FLAG tag in red (IRDye 680RD Donkey anti-mouse); middle panel: Hartmani NP in green (IRDye 800CW Donkey anti-rabbit); right panel: merged image. Immunodetection was performed using the Odyssey Infrared Imaging System (LICOR, Biosciences) providing also the molecular marker (Precision Plus Protein Dual Color Standards, Bio-Rad) used. See also Supplementary Figure 3.

### Cell fractionation supports NP’s mitochondrial localization

To confirm mitochondrial localization of arenavirus NP, we performed a kit-based subcellular fractionation of cells infected with the “equimolar” mix of UGV-1, UHV-1 and ABV-1. In the immunoblotting-based analyses, tubulin served as marker for the cytosolic fraction, and MFN2, mitochondrial ribosomal protein S35 (MRPS35), voltage-dependent anion-channel (VDAC), and cytochrome c oxidase subunit IV (COX IV) as mitochondrial markers. A reference for loading is provided by Ponceau S staining (Supplementary Fig. 4a). At one dpi, the mitochondrial fraction of reptarenavirus infected V/4 Br cells contained a low amount of NP, as expected for the early phase of infection; it had increased by two dpi (Fig. 4a). Infection of *Boa constrictor* I/1 Ki and V/4 Lu cells yielded similar results (Supplementary Figs. 5a, 6a). In parallel, we isolated mitochondria from V/4 Br cells infected with the same virus mix, using a different method for obtaining mitochondria preparations. Cells and fractions were collected at one, two and four dpi and analyzed by immunoblotting with tubulin as cytosolic, and MFN2, VDAC and COX IV as mitochondrial markers (Ponceau S staining of the membrane, Supplementary Fig. 4b). Again, reptarenavirus NP was present in the mitochondrial fraction of infected cells from one dpi onwards (Fig. 4b). Infection and fractionation of I/1 Ki cells produced similar results (Supplementary Figs. 5b, 6b).

**Figure 4.**
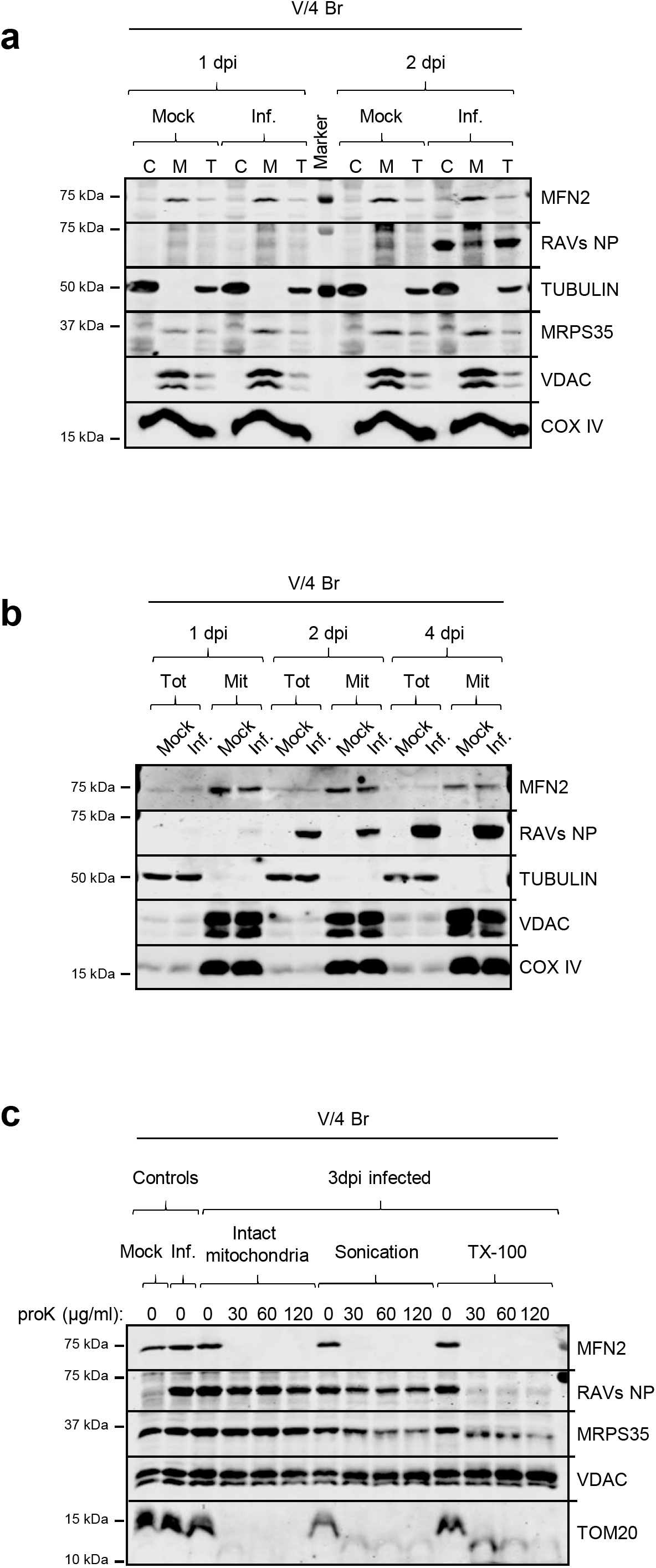
Subcellular and submitochondrial analyses of reptarenaviral NP in infected boid cells. **(a)** Subcellular fractionation analyses of *Boa constrictor* V/4 Br cells infected with the “equimolar” reptarenavirus mix of UGV-1, UHV-1 and ABV-1, at one and two dpi. Uninfected (Mock) samples provide the negative controls. Cytosolic (C) and mitochondrial (M) fractions and total cellular extracts (T) were obtained from a standard cell fractionation procedure and separated in SDS-PAGE gels, followed by immunoblotting. Antibodies detecting tubulin (a cytosolic marker), COX IV, VDAC, MRPS35 and MFN2 (mitochondrial markers), and reptarenaviral NP were used. **(b)** Immunoblot analyses of whole-cell lysates (Tot) and mitochondrial preparations (Mit) obtained from *Boa constrictor* V/4 Br cells inoculated with the “equimolar” reptarenavirus mix of UGV-1, UHV-1 and ABV-1 and analyzed at one, two and four dpi. Uninfected (Mock) samples provide the negative controls. 25 μg protein samples were separated on SDS-PAGE followed by immunoblotting to detect tubulin (a cytosolic marker), COX IV, VDAC, and MFN2 (mitochondrial markers) and reptarenaviral NP. **(c)** Submitochondrial localization assay, determined by protease accessibility. Mitochondria were isolated at three dpi from *Boa constrictor* V/4 Br cells inoculated with the “equimolar” reptarenaviral mix of UGV-1, UHV-1 and ABV-1 and either treated directly with Proteinase K (proK) at 30, 60 or 120 μg/ml, or subjected to sonication or Triton X-100 (TX-100) lysis first, and then treated with proK. For each condition, a proK-untreated sample is provided as control. An uninfected (Mock) and a reptarenavirus-infected mitochondrial sample at three dpi are present respectively as negative and positive controls for the anti-UHV NP antibody used to detect the reptarenaviral NPs. 25 μg mitochondrial samples were separated through standard SDS-PAGE followed by immunoblotting to detect TOM20 and MFN2 (markers of the outer mitochondrial membrane), MRPS35 (marker of mitochondrial matrix), VDAC (loading control) and reptarenaviral NP. VDAC, embedded in the outer mitochondrial membrane, is not affected by proK treatment and thus provides an internal reference for loading. Immunodetections were performed using the Odyssey Infrared Imaging System (LICOR, Biosciences), showing also the molecular weight marker (Precision Plus Protein Dual Color Standards, Bio-Rad) used. See also Supplementary Figures 4-6.

Both subcellular fractionation and mitochondria isolation methods rely on differential centrifugations, which could lead to co-purification of IBs and mitochondria. To exclude the latter and to confirm that a fraction of NP indeed resides inside mitochondria, we performed protease-based submitochondrial localization analyses^48^. We compared the effect of increasing proteinase K (proK) concentrations on intact, sonicated, and Triton X-100 (TX-100) treated mitochondria, freshly isolated at three dpi from V/4 Br cells infected with the aforementioned viral mix, and analyzed the samples by immunoblotting. While proteins localized at the cytosolic side of the mitochondrial outer membrane are supposedly accessible for proK under all test conditions, the proteins in the intermembrane space, on the inner membrane and in the matrix are inaccessible to proK in intact mitochondria. Sonication, and TX-100 treatment respectively served as mild and robust way of disrupting the mitochondrial membranes to render all mitochondrial proteins susceptible to proK degradation. The results show degradation of the mitochondrial outer membrane translocase 20 (TOM20) and MFN2, proteins located in the outer membrane, under all conditions (Fig. 4c). A lack of competing substrates in intact mitochondria could explain the slower degradation of TOM20 following sonication and TX-100 treatment. VDAC, firmly embedded in the outer mitochondrial membrane, appeared inaccessible to proK under all test conditions (Fig. 4c), and we thus used it as an internal loading control. A Ponceau S staining of the membrane is also shown as evidence of protein degradation (Supplementary Fig. 4c). Unfortunately, we did not find a suitable marker protein for the intermembrane space of snake mitochondria. MRPS35, which resides in the mitochondrial matrix, remained intact upon proK treatment of intact mitochondria and degraded following sonication and TX-100 treatment (Fig. 4c). ProK treatment of intact mitochondria caused degradation of a small amount of reptarenavirus NP, suggesting that some cytosolic NP co-purifies with the mitochondria. However, sonication resulted in slight NP degradation, while almost complete loss of NP occurred following TX-100 treatment (Fig. 4c). The results indicate similar degradation patterns for MRPS35 and NP, adding to the evidence that NP localizes in the mitochondrial matrix.

### NPs within mammalian mitochondria contain the predicted MTS

To study if the NPs enter the mitochondria of mammalian cells and whether the putative MTS would be cleaved in them, we transfected African green monkey kidney (Vero E6) cells with the following constructs: wtUGV1-NP-FLAG, mutUGV1-NP-FLAG, wtJUNV-NP-HA, mutJUNV-NP-HA, UHV1-NP-FLAG, LCMV-NP-HA, and HISV1-NP-FLAG. Because we earlier demonstrated that reptarenaviruses replicate at 30°C rather than at 37°C^49^, we performed the experiments at both temperatures and used immunoblotting and IF for analyses.

Immunoblotting of whole-cell lysates at three dpt using anti-FLAG or anti-HA antibodies showed that all wt and mutNP constructs produced a protein of approximately 63-68 kDa in molecular weight at both temperatures (Figs. 5a,b, 6a). For HISV-1 NP we confirmed the result also by using the anti-Hartmani NP antibody (Fig. 5b). Re-probing of the membranes with anti-MTS antiserum showed that the putative MTSs remained uncleaved (Figs. 5a, 6a), as observed for the reptilian cells (Fig. 3a-c). Curiously, the expression of both wt and mutUGV1 NPs as well as LCMV NP was stronger at 30°C than at 37°C (Figs. 5a, 6a). The phenomenon was most evident for HISV NP which was barely detectable after incubation at 37°C (Fig. 5b).

**Figure 5.**
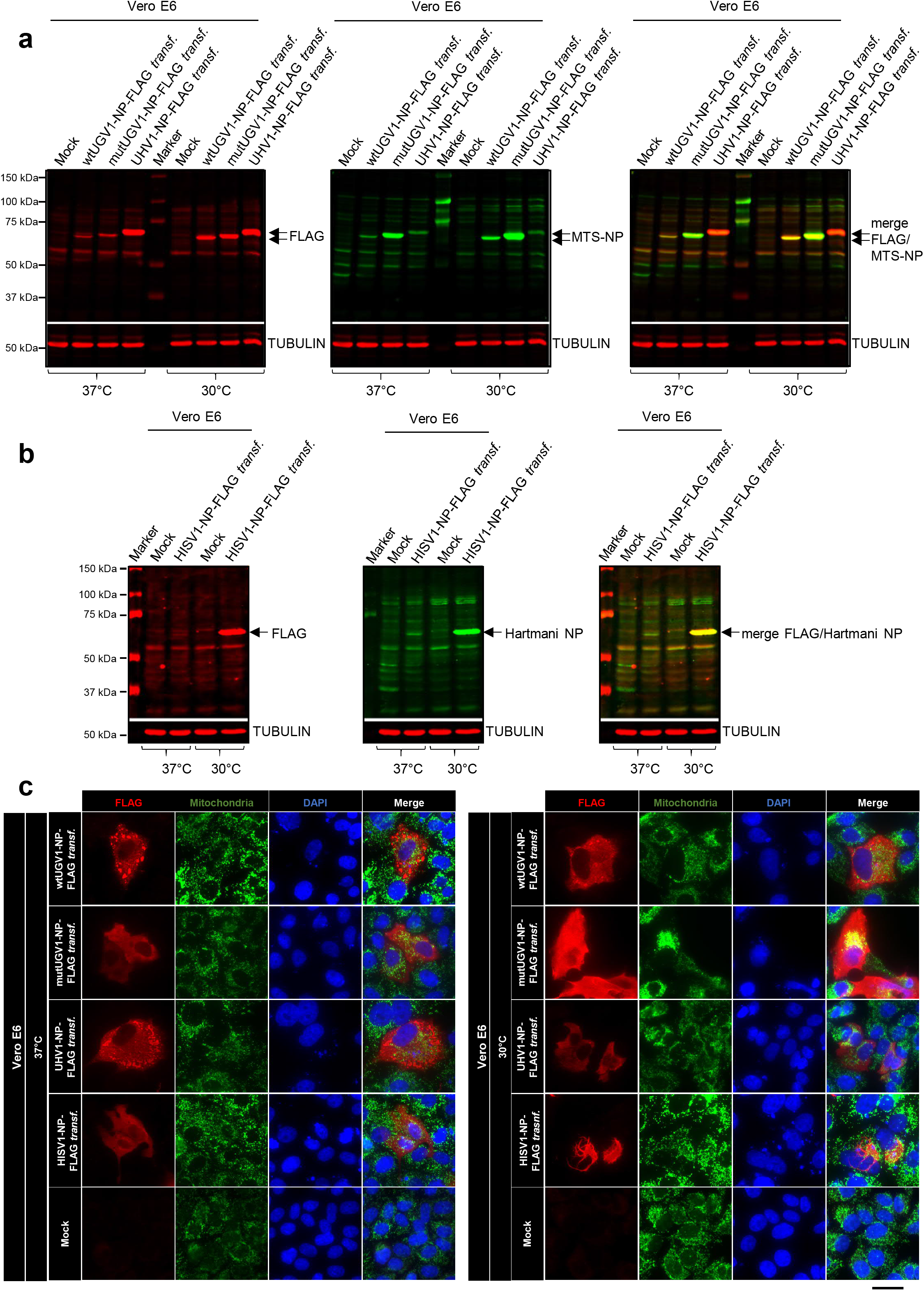
Immunoblotting and IF studies on reptarenaviral or hartmanivirus NPs in transfected mammalian cells. **(a,b)** Immunoblot analyses of whole-cell lysates obtained from monkey Vero E6 cells transfected with a construct expressing either wt or mutUGV1-NP-FLAG, or UHV1-NP-FLAG **(a)** and HISV1-NP-FLAG **(b)**, at three dpt after incubation at either 37°C or 30°C. Non-transfected (Mock) samples are provided as negative controls. 40 μg protein samples were loaded on standard SDS-PAGE gels, followed by immunoblotting analyses. Tubulin (in red) was used as a reference for loading. Reptarenavirus and hartmanivirus NPs (65-68 kDa) are indicated (black arrows). **(a)** Left panel: FLAG tag in red (IRDye 680RD Donkey anti-mouse); middle panel: MTS-NP in green (IRDye 800CW Donkey anti-rabbit); right panel: merged image. **(b)** Left panel: FLAG tag in red (IRDye 680RD Donkey anti-mouse); middle panel: Hartmani NP in green (IRDye 800CW Donkey anti-rabbit); right panel: merged image. Immunodetection was performed using the Odyssey Infrared Imaging System (LICOR, Biosciences) providing also the molecular marker (Precision Plus Protein Dual Color Standards, Bio-Rad) used. **(c)** Double IF images of monkey Vero E6 cells transfected with a construct expressing either wt or mutUGV1-NP-FLAG, UHV1-NP-FLAG or HISV1-NP-FLAG at three dpt, after incubation at either 37°C (left panel) or 30°C (right panel). Non-transfected (Mock) cells served as controls. The panels from left: FLAG tag in red (AlexaFluor 594 goat anti-rabbit), mitochondrial marker (mtCO2) in green (AlexaFluor 488 goat anti-mouse), nuclei in blue (DAPI), and a merged image. Scale bar: 200 μm. See also Supplementary Figures 7 and 8.

**Figure 6.**
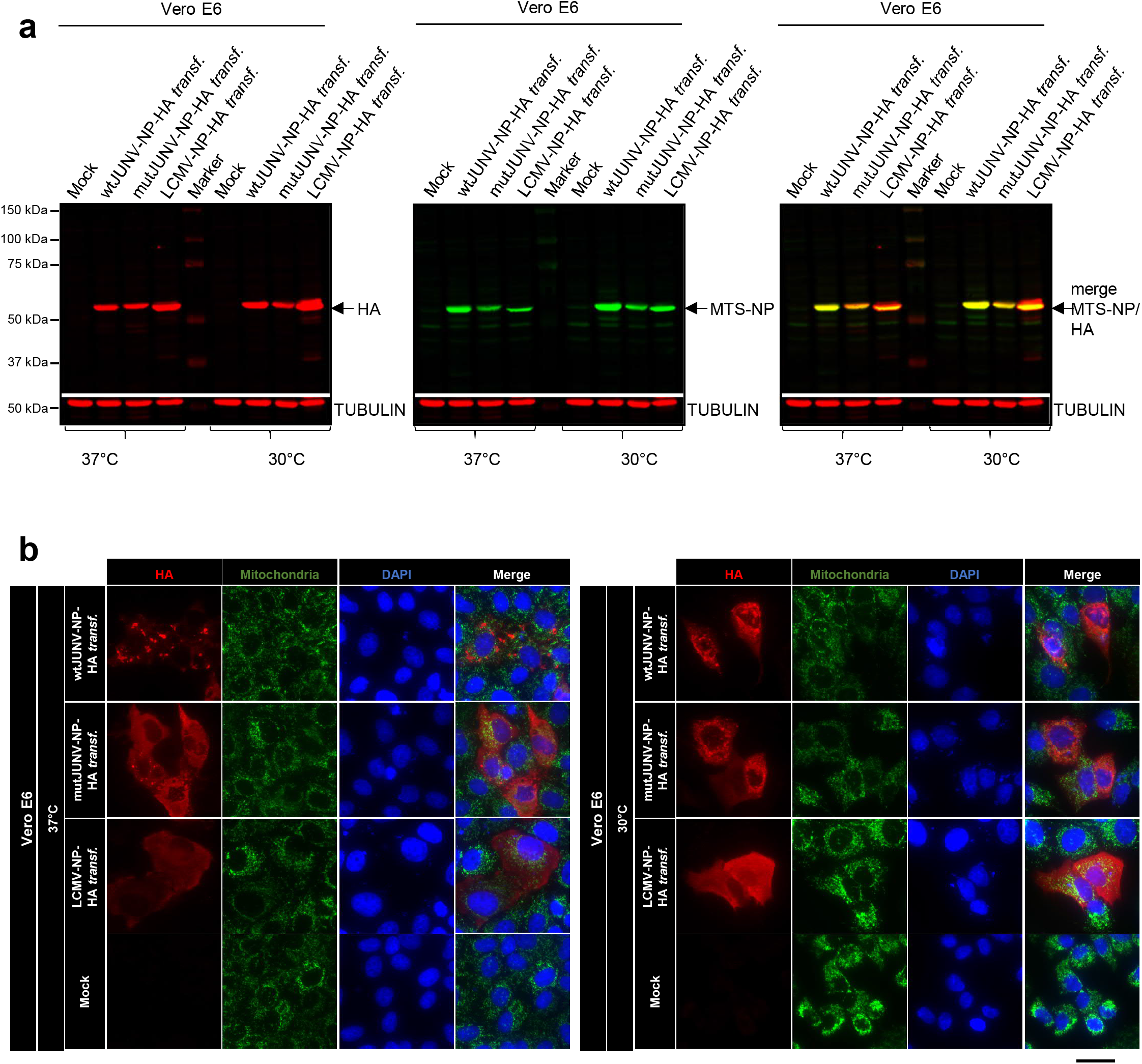
Immunoblotting and IF studies on mammarenaviral NPs in transfected mammalian cells. **(a)** Immunoblot analyses of whole-cell lysates obtained from monkey Vero E6 cells transfected with a construct expressing either wt or mutJUNV-NP-HA, or LCMV-NP-HA at three dpt, after incubation at either 37°C or 30°C. Non-transfected (Mock) samples are provided as negative controls. 40 μg protein samples were loaded on standard SDS-PAGE gels followed by immunoblotting. Tubulin (in red) was used as a reference for loading. Mammarenavirus NPs (63-65 kDa) are indicated (black arrows). Left panel: HA tag in red (IRDye 680RD Donkey anti-mouse); middle panel: MTS-NP in green (IRDye 800CW Donkey anti-rabbit); right panel: merged image. Immunodetection was performed using the Odyssey Infrared Imaging System (LICOR, Biosciences) providing also the molecular marker (Precision Plus Protein Dual Color Standards, Bio-Rad) used. **(b)** Double IF images of monkey Vero E6 cells transfected with a construct expressing either wt or mutJUNV-NP-HA, or LCMV-NP-HA at three dpt, after incubation at either 37°C (left panel) or 30°C (right panel). Non-transfected (Mock) cells served as controls. The panels from left: HA tag in red (AlexaFluor 594 goat anti-rabbit), mitochondrial marker (mtCO2) in green (AlexaFluor 488 goat anti-mouse), nuclei in blue (DAPI), and a merged image. Scale bar: 200 μm. See also Supplementary Figure 9.

At three dpt, after incubation at 37°C, IF analyses showed that wtUGV1, UHV1 and wtJUNV NP yielded a punctate staining pattern, while both mutUGV1 and mutJUNV NPs as well as LCMV NP yielded a diffuse staining pattern (Figs. 5c, 6b), as previously described for reptilian cells incubated at 30°C (Fig. 2c-f). The IF analysis for HISV-1 NP concurred with the immunoblot, a weak and diffuse staining pattern was seen in cells incubated at 37°C whereas cells kept at 30°C presented more intense staining with tubular structures around the perinuclear area (Fig. 5c). At 30°C, while wtUGV1, wtJUNV and UHV1 NP showed mainly a punctate staining pattern, and mutUGV1, mutJUNV and LCMV NP a diffuse pattern, wtNPs also yielded some diffuse staining and mutUGV1 and mutJUNV NP formed occasional aggregates (Figs. 5c, 6b). For the reptarenavirus NPs (wt and mutUGV1, UHV1 NP), in Vero E6 cells the pan-reptarenavirus NP antiserum produced similar staining patterns as the anti-FLAG antiserum (Supplementary Fig. 7a,b). Using the anti-MTS antibody on cells incubated at either temperature yielded a prominent diffuse staining for the mutNPs and no signal for the wtNPs including LCMV NP (Supplementary Figs. 8a,b, 9a,b), similar to what we observed in reptilian cells (Supplementary Fig. 2a-d). Interestingly, staining for arenavirus NPs and the mitochondrial marker mtCO2 showed a similar association in mammalian cells as in the reptilian cells: while there was no clear evidence of co-localization, some wtNP aggregates were apparently associated with the mitochondrial network (Figs. 5c, 6b). However, after incubation at 30°C the mitochondrial network appeared condensed, which could relate to the fact that mammalian cells are adapted to 37°C. The suboptimal temperature could explain the slight differences observed between the distribution patterns of the analyzed arenavirus NPs in reptilian *vs* mammalian cells incubated at 30°C (Figs. 2c-f, 5c, 6b). To conclude, the results in mammalian cells concur with those obtained in reptilian cells.

### NP mitochondrial translocation does not occur *in vitro*

To understand the mechanism behind the mitochondrial translocation of arenavirus NPs, we performed an *in vitro* mitochondrial import assay utilizing mitochondria freshly isolated from I/1 Ki, *Python regius* heart (VI/1 Hz) and Vero E6 cells. First, we compared the ability of I/1 Ki, VI/1 Hz and Vero E6 cells to mediate the import of an *in vitro* translated chimeric mitochondrial fusion protein, Su9-DHFR (MTS of Subunit 9 of mitochondrial ATPase, Su9, of *Neurospora crassa* fused with the dihydrofolate reductase, DHFR, of *Mus musculus*^50,51^). The subsequent analysis identified the precursor, intermediate and mature mitochondria-imported forms of Su9-DHFR in the samples generated with mitochondria isolated from all tested cell lines (Fig. 7a-c). Carbonyl cyanide m-chlorophenyl hydrazine (CCCP) treatment induces loss of mitochondrial membrane potential, thus significantly decreasing or abolishing MTS-mediated mitochondrial import^52^, and only the Su9-DHFR precursor and intermediate forms could be detected following the treatment (Fig. 7a-c). ProK treatment prior to electrophoresis led to loss of the Su9-DHFR precursor and intermediate forms in samples with intact (CCCP-untreated) mitochondria, while the mature form was protected from degradation as it was located within the mitochondria; all bands were lost following proK treatment of CCCP-treated samples (Fig. 7a-c). The Su9-DHFR translocation occurred at both 30°C and 37°C with similar efficacy (Fig. 7a-c). These results indicate that mitochondria isolated from cultured snake cells remain intact, and that the mitochondrial import system functions in a similar way in reptilian and mammalian cells.

Next we assessed the mitochondrial import of wtUGV1-NP, mutUGV1-NP, UHV-1, HISV-1, JUNV and LCMV NP and produced the proteins via *in vitro* translation under the control of either the SP6 or T7 promoter. The result indicates that arenavirus NPs cannot be translocated into mitochondria *in vitro*, since none of the proteins remained detectable following proK treatment (Fig. 7d-h; Supplementary Fig. 10a-d).

**Figure 7.**
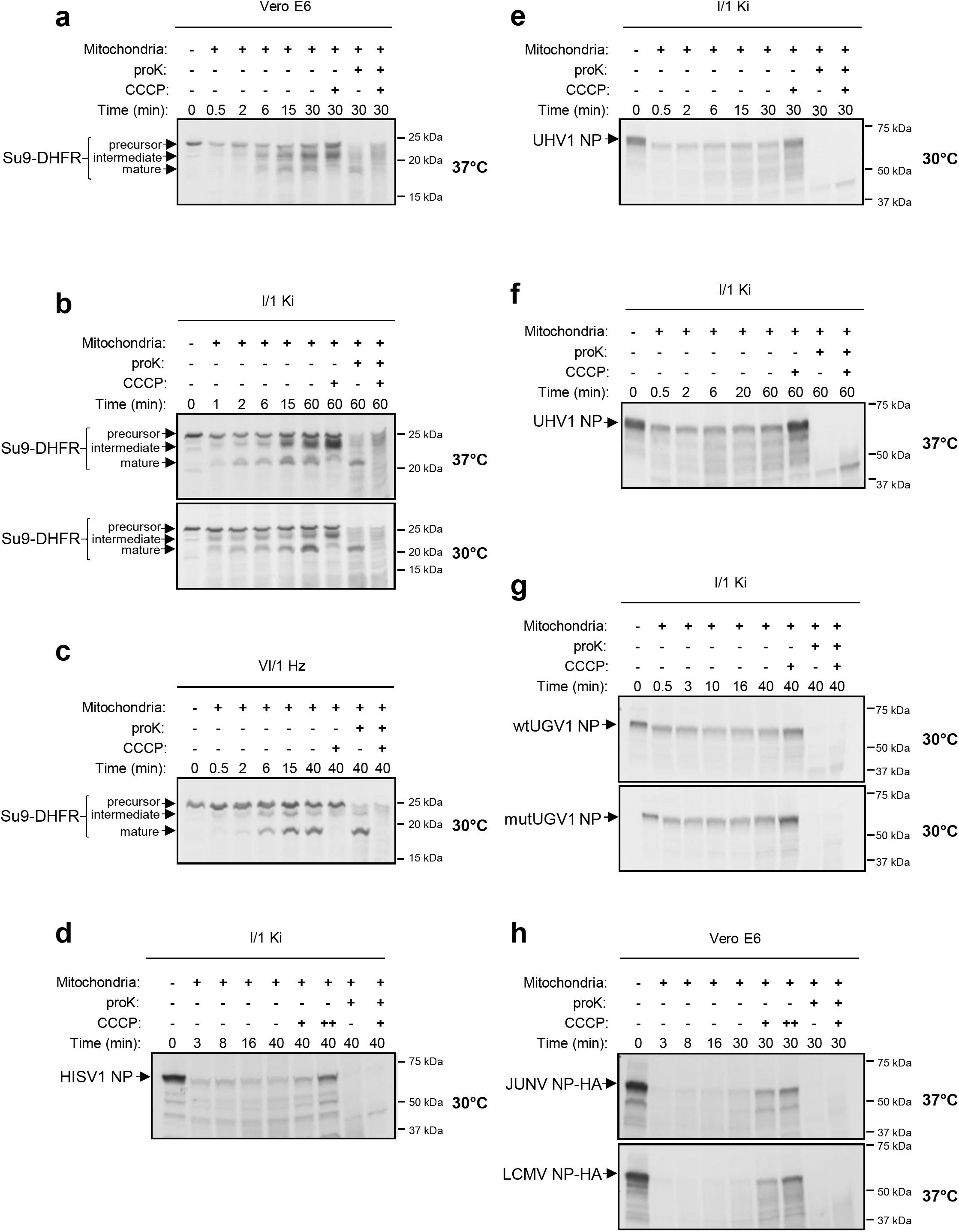
*In vitro* import into mitochondria of arenaviral NPs. **(a-c)** The *in vitro* import into mitochondria was determined for a known chimeric mitochondrial protein, Su9-dihydrofolate-reductase (DHFR), composed of the MTS of Subunit 9 of *N. crassa* ATPase (Su9) fused to *M. musculus* DHFR, and used as positive control for the assay. The fusion protein was synthesized using [^35^S]-methionine in a rabbit reticulocyte lysate, from its cDNA sequence cloned in a pGEM4Z vector under the control of the SP6 RNA promoter. The hybrid protein was imported into freshly isolated mitochondria of monkey Vero E6 cells, at 37°C **(a)**, *Boa constrictor* kidney (I/1 Ki) cells, at both 37°C and 30°C **(b)**, and *Python regius* heart (VI/1 Hz) cells, at 30°C **(c)**, as indicated by the presence at different time points of three distinct translocation forms: precursor, intermediate and mature forms (black arrows). **(d-h)** The *in vitro* translocation into freshly isolated *Boa constrictor* I/1 Ki mitochondria was assessed for HISV-1 NP, at 30°C **(d)**, UHV-1 NP, at 30°C **(e)** and 37°C **(f)**, wt and mutUGV-1 NPs, at 30°C **(g)** and HA-tagged JUNV and LCMV NPs, at 37°C **(h)**. Radiolabelled NPs were *in vitro* synthesized using [^35^S]-methionine in a rabbit reticulocyte lysate, from their cDNA sequence cloned into pGEM4Z **(d-g)** or pCR4Blunt-TOPO **(h)** vectors under the control of the SP6 **(d-g)** or T7 **(h)** promoter. Protein signals were determined through autoradiographic detection. CCCP: mitochondrial protein import blocker by inducing mitochondrial membrane potential dissipation. Proteinase K (proK): leading to degradation of non-imported proteins. See also Supplementary Figure 10.

### Immune EM confirms mitochondrial IBs as arenavirus NP

Having collected evidence of mitochondrial localization of arenavirus NPs by molecular biology techniques, and given the apparent ultrastructural similarity of the intramitochondrial electron-dense structures with the reptarenavirus-induced IBs, immune EM was attempted to validate the findings. Indeed, immunogold labeling of UGV-1 infected I/1 Ki cells at three dpi demonstrated the presence of NP, both in the cytoplasm and within the mitochondria (Fig. 8a). UHV1-NP-FLAG transfected I/1 Ki cells at three dpt yielded even more prominent NP staining within mitochondria (Fig. 8b-f), which displayed both electron-dense and electron-lucent IBs and smaller IBs (Fig. 8b-f), confirming the observations made in infected cells (Figs. 1a-c, 8a).

**Figure 8.**
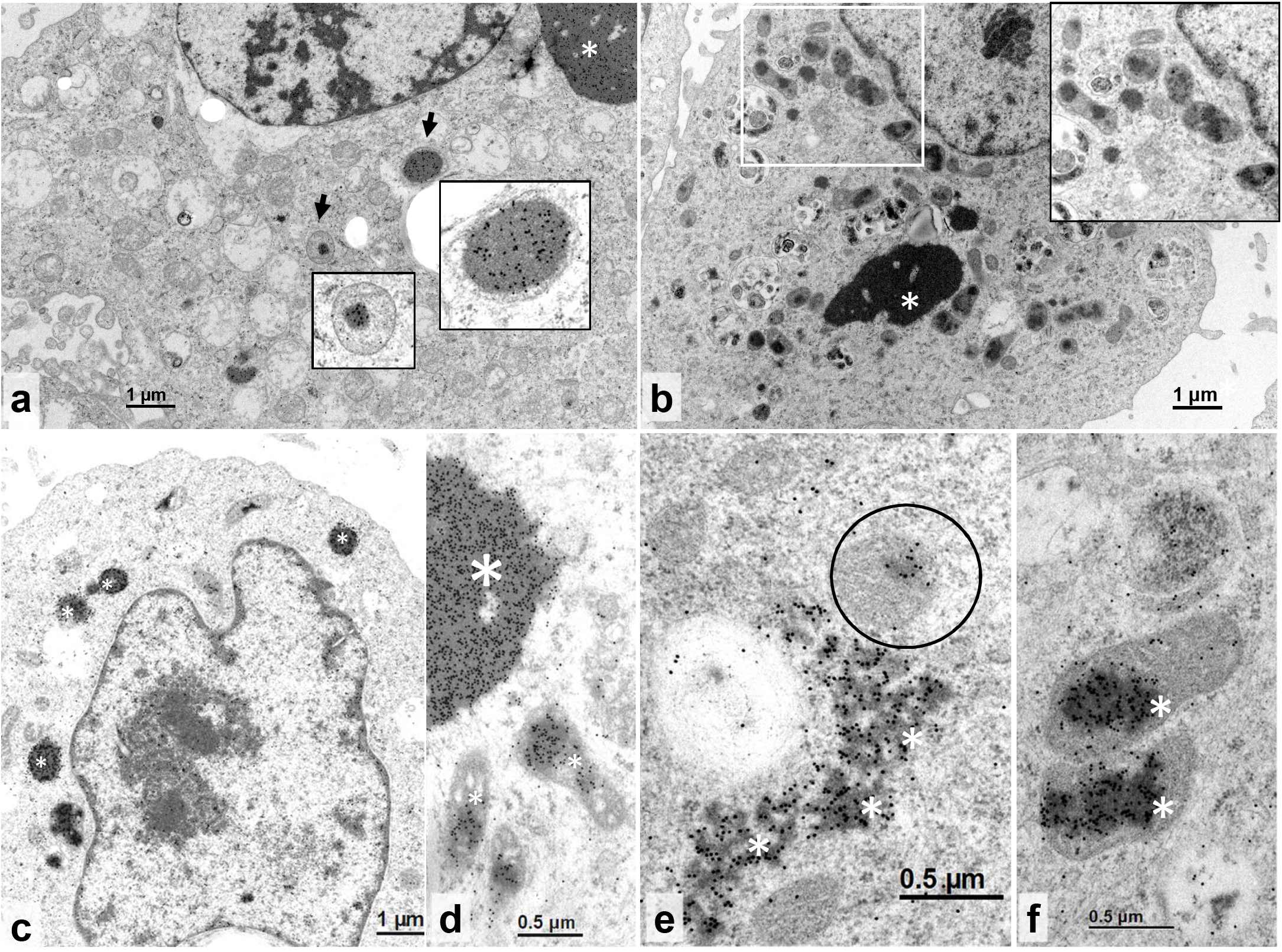
TEM and immune EM on reptarenavirus-infected or NP-transfected boid cell lines. **(a)** Permanent cell culture derived from *B. constrictor* kidney (I/1 Ki), infected with UGV-1, at three dpi. Immunogold labelling of the reptarenaviral NP. Large NP-positive cytoplasmic IB (asterisk) and several NP-positive IBs within the matrix of mitochondria (arrows). Inserts: higher magnification of the areas indicated by the arrows. **(b-f)** Permanent cell culture derived from *B. constrictor* kidney (I/1 Ki), transfected with UHV1-NP-FLAG, at three dpt. **(b)** Cell with large electron-dense IB (asterisk) and multiple mitochondria with IB formation (highlighted by a white rectangle). Insert: higher magnification of the area depicted in the rectangle. **(c-f)** Immunogold labelling of reptarenaviral NP. **(c)** Positive reaction in small electron-dense cytoplasmic IBs (asterisks). **(d)** Large electron-dense IB with positive reaction (larger asterisk) and individual mitochondria with positive IBs within the matrix (smaller asterisks). **(e)** Irregular shaped, more electron-lucent, presumably earlier cytoplasmic IB (asterisk) and single mitochondrion with positive reaction within the matrix (circle). **(f)** Mitochondria with positive IBs within the matrix (asterisks) at higher magnification.

Finally, we wanted to determine whether the mitochondrial IB localization observed *in vitro* would also occur *in vivo* and examined the brain of *Boa constrictor* snakes with confirmed BIBD. Both the electron-dense, variably sized round IBs with a smooth outline and the irregularly shaped, less electron-dense IBs with irregular borders that we saw *in vitro* (Figs 1a-c, 8a-f) were also present in neurons in the brain (Fig. 9a-f). Mitochondria that exhibited the less electron-dense IBs were often swollen, partly vacuolated and disrupted, with a granular, electron-dense matrix and indistinct, possibly ruptured outer membrane (Fig. 9a-d). Both types of IBs as well as occasional mitochondria contained viral NP, as shown by the immunogold labelling (Fig. 9e-f), thus indicating that the *in vitro* findings are translatable to the *in vivo* situation during natural infection.

**Figure 9.**
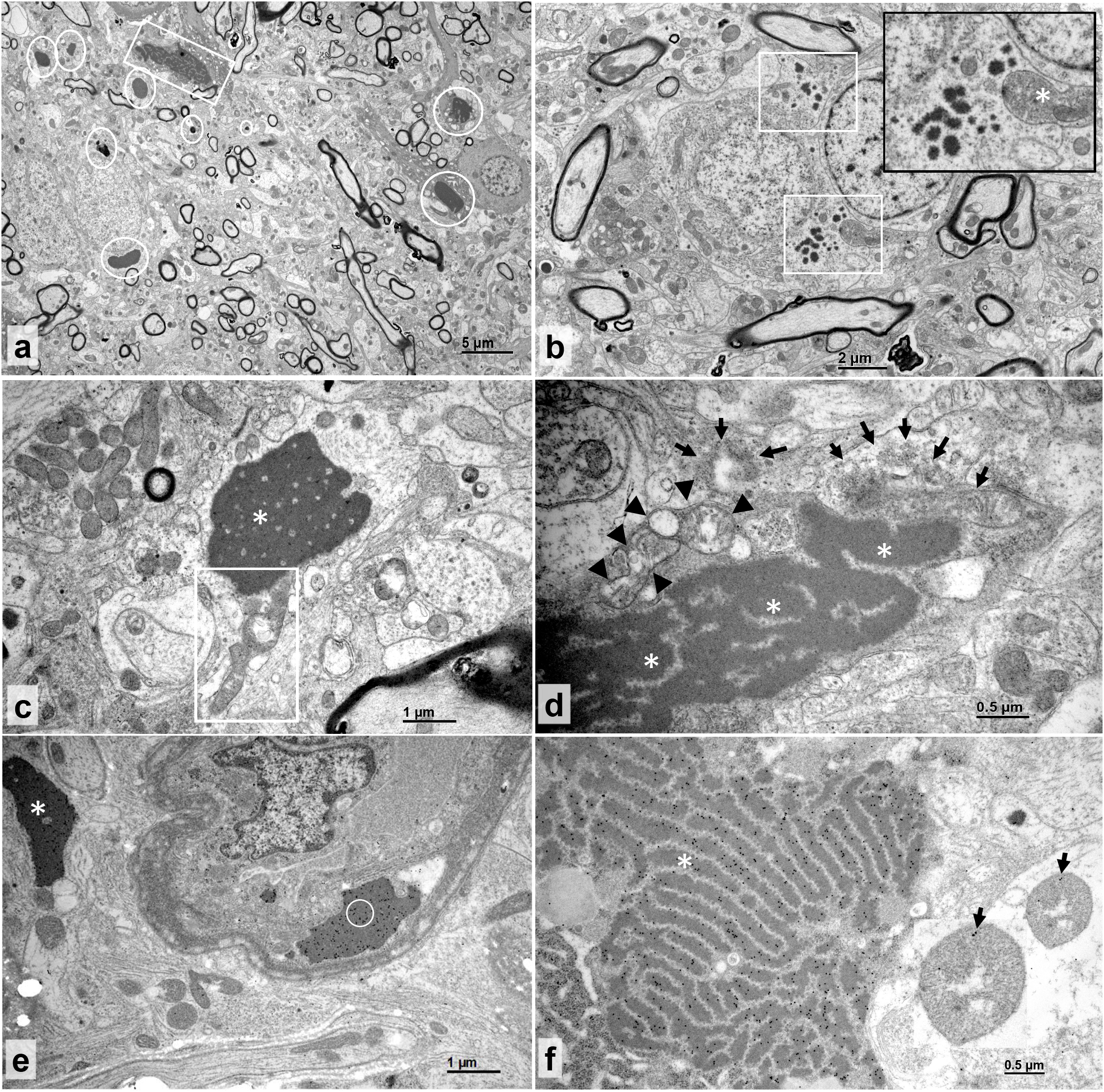
TEM and immune EM of a BIBD-positive *B. constrictor* brain. **(a-d)** TEM, neurons. **(a)** Numerous <1-3.5 μm sized, round, smooth edged electron-dense cytoplasmic IBs (circles) and one larger, less electron-dense, irregularly shaped IB with more coarse margins (square). **(b)** Neurons with small, 0.1-3 μm sized electron-dense cytoplasmic IBs (white squares). Insert: higher magnification depicting irregularly shaped IB borders and a swollen mitochondrion with disorganized coarse electron-dense matrix (asterisk). **(c)** Mitochondrion with vacuolated dissolved matrix (rectangle), adjacent to a larger IB (asterisk). **(d)** Large cytoplasmic IB (asterisks) with adjacent vacuolated mitochondria (arrowheads) and mitochondria with granular, dissolved matrix (arrows). **(e-f)** Immunogold labelling. **(e)** NP-positive electron-dense cytoplasmic IBs (asterisk) within a neuron adjacent to a small vessel with an IB (circle) in an endothelial cell. **(f)** NP-positive, less electron-dense irregular cytoplasmic IB (asterisk) and focal positive reaction within mitochondria (arrows).

## DISCUSSION

Reptarenavirus NP is intimately linked to BIBD since it is the main component of the IBs that are pathognomonic for the disease and manifest in numerous cell types. Ultrastructural studies on reptarenavirus infected cells revealed IBs within mitochondria; this prompted us to study the phenomenon and mechanisms of the potential mitochondrial transport of arenavirus NPs in detail. Software prediction tools identified a putative MTS in the N-terminus of rept- and mammarenavirus but not in hartmanivirus NPs. Mutagenesis studies indicated that the putative MTS contributes to aggregate and, possibly, IB formation of rept- and mammarenavirus NPs. However, cell fractionation, protein-processing analysis including immunoblots using anti-MTS antibody, and *in vitro* import assays indicated that the putative MTS does not mediate mitochondrial translocation. Analysis of mitochondria isolated from both infected and transfected cells demonstrated that the NPs of reptarena-, hartmani-, and mammarenavirus enter the mitochondria, suggesting that the feature is common among arenaviruses. The fact that hartmanivirus NPs lack the putative MTS, combined with experimental evidence from mutagenesis analyses on the putative MTSs, suggests that the mitochondria translocation occurs via an internal translocation signal (ITS). Finally, we employed immune EM to demonstrate that the IBs found within mitochondria indeed represent reptarenavirus NP. Strikingly, by studying the brain of snakes with BIBD by TEM, we could demonstrate that mitochondrial translocation of reptarenavirus NP occurs also *in vivo*.

Our first idea, after identifying NP inside the mitochondria of reptarenavirus infected cells, was to confirm that mitochondrial ribosomes cannot mediate its translation. After confirming that mitochondria codons would not produce full-length protein, we hypothesized that the NP must contain an MTS. The analyses involving two software tools identified a putative MTS in the NPs of several rept- and mammarenavirus but not in those of hartmaniviruses. To our surprise, experiments with wt and mut forms of rept- and mammarenavirus NPs did not result in cleavage, although this commonly occurs following MTS-mediated mitochondrial translocation^46^. However, mitochondrial import can occur without MTS cleavage, as shown for, e.g., the 10 aa MTS sequence of the human T-cell leukemia virus type 1 (HTLV-1) p13II protein which remains intact during import^53^. In addition, our results showed that mutations altering the positive charges in the putative MTS did not affect mitochondrial localization, even though they reduced the software-predicted mitochondrial translocation potential. However, not all proteins enter the mitochondria with the help of the classical N-terminal MTS; there might be additional internal cryptic elements within the NP that contribute to the mitochondrial localization. For instance, Hepatitis B virus (HBV) Pol protein localizes to both cytosol and mitochondria, and it can enter the mitochondria even after removal of its intrinsic MTS, suggesting the presence of multiple MTSs^54^.

We set up a mitochondrial import assay with reptilian mitochondria, and used a control protein to verify the functionality before studying arenavirus NP translocation. The *in vitro* assay could not induce mitochondrial translocation of NP. Co-translational mitochondrial import, in which only the newly translated protein originating from ribosomes close to mitochondria becomes translocated^55^, might be one of the factors contributing to NP translocation. In the *in vitro* assay, the protein synthesis occurred prior to addition of isolated mitochondria, and the procedure did apparently not preserve the conditions favoring mitochondrial import. It is possible that the mitochondrial import of NP relies on factors absent in the *in vitro* assay or involves non-classical ways, e.g. by translocases. For instance, the transcription factor p53, an apoptosis inducer, normally resides in the cytosol and/or nucleus but under certain conditions targets the mitochondria via binding to the MTS-bearing Tid1 factor^56,57^. Another possible explanation can lie in a phenomenon known as “eclipsed distribution”, where the higher amount of a protein in a cell compartment obscures its detection in another compartment^58^. This would suggest the possibility that only a small subpopulation of NP, hardly detectable, would enter the mitochondria while the rest would localize elsewhere. Therefore, it would be possible that the amount translocated to mitochondria remained below the detection limit in our *in vitro* assay.

Dual protein localization is a dynamic process responding to cellular conditions or aiming at rebalancing the distribution of protein subpopulations between cellular compartments. The subcellular localization of dually localized proteins depends e.g. on the targeting signal, folding, proteolytic cleavage, binding to other proteins, and on post-translational modifications^57^. For example, Human Herpesvirus-8 (Kaposi’s sarcoma-associated herpesvirus) harbors two anti-apoptotic proteins that have multiple localizations: the K7 protein localizes to ER, nucleus and mitochondria, and the KS-Bcl2 protein either to mitochondria or nucleus. Similarly, the hepatitis C virus proteins Core, p7, and NS3/4A localize to mitochondria and/or mitochondria-associated membranes in addition to the ER, and the Core protein also to lipid droplets. Also, the human papilloma virus E1^E4 protein is distributed between mitochondria and the cytokeratin network, and the E2 protein among nucleus, cytosol and mitochondria^59^. The software predictions for arenavirus NP would fit its dual localization to cytosol and mitochondria. The dual localization of (rept)arenavirus NPs might e.g. be driven by the oligomerization status/quaternary structure of the proteins, which implies that the translocation signal (terminal or internal) might be accessible only in the unfolded state, and possibly also in the monomeric or oligomeric form of the protein. Also, the folding state of the NP could induce its reverse translocation back to the cytoplasm during the import process, a phenomenon that has been suggested to determine, for example, the subcellular localization of the *S. cerevisiase* fumarase^60^. Alternatively, RNA binding could have an impact on the mitochondrial translocation of NPs, e.g. only NPs free from RNA would enter the mitochondria.

Both mamm- and reptarenavirus infections are non-cytopathic in cell culture^31,49,61^, while hartmanivirus infection appears to be cytopathogenic^16^. Mammarenaviruses NPs are involved in suppression of IFN signaling^17–24^, and the secondary infections that often cause the death of snakes with BIBD could be the result of similar immunosuppressive functions of reptarenavirus NP. Supporting the hypothesis are the facts that snakes with BIBD exhibit low amounts of antibodies against reptarenavirus NPs and that white blood cells are among the cell types where IBs are particularly prominent^31,38,40^. While the immunosuppression by arenaviruses is mechanistically linked to the ability of NP to prevent IFN signaling^17–24^, mitochondrial targeting could further contribute to the dampening of the innate immune response. For example, the severe acute respiratory syndrome coronavirus open-reading frame 9b protein localizes to mitochondria and causes MAVS signalosome degradation^62^. The PB1-F2 protein of influenza A virus localizes in the mitochondrial intermembrane space, where it suppresses the immune response and induces activation of the nucleotide-binding domain (NOD)-like receptor protein 3 (NLRP3) inflammasome by altering the mitochondrial membrane potential^63^. Hepatitis C virus NS3/4A protease on the other hand localizes to the outer mitochondrial membrane and mediates MAVS cleavage, thus suppressing the downstream IFN signaling^64,65^.

We herein provided *in vitro* and *in vivo* evidence showing that mitochondrial localization is a common feature of arenavirus NPs. Transfection resulted in mitochondrial localization of rept-(UGV-1 and UHV-1) and mammarenavirus (JUNV and LCMV) as well as hartmanivirus (HISV-1) NP, and mutations to the putative MTSs did not prevent the localization but interfered with NP aggregation. Our observations further suggest that arenavirus NPs could either possess cryptic or internal mitochondrial translocation signals that are not recognized by bioinformatic softwares, or that the NPs may employ alternative strategies for mitochondrial localization. The presence of arenavirus NP within mitochondria could indicate a previously unknown mechanism to suppress the innate immune system. We propose that by targeting mitochondria (rept)arenaviruses could (*i*) reduce their cytoplasmic presence to escape pathogen-recognition factors and evade the innate immune response; (*ii*) affect mitochondrial functions in immune response by affecting MAVS or inducing mitophagy to avoid apoptosis; (*iii*) induce mitochondrial biogenesis to control the metabolic and redox state of the cell. All these hypotheses on the possible role of mitochondrial localization of reptarenaviral NP require further investigations.

## MATERIAL AND METHODS

### Cell lines and viruses

The study made use of the African green monkey kidney, Vero E6 (American Type Culture Collection [ATCC]) cell line, and permanent tissue cell cultures derived from *Boa constrictor* brain (V/4 Br), kidney (I/1 Ki)^31^, lung (V/4 Lu) and liver (V/1 Liv), and *Python regius* heart (VI/1 Hz). The cells were maintained in Minimum Essential media (MEM, Gibco), containing 10% fetal bovine serum (FBS, Biochrom), 10% tryptose phosphate broth (TPB, Difco), 6 mM Hepes (Biochrom), 2 mM L-alanyl-L-glutamine (Biochrom) and 50 μg/ml Gentamicin (Gibco). The snake cells were maintained at 30°C and mammalian cells at 30°C or 37°C, with 5% CO_2_.

For infections, the single University of Giessen virus 1 (UGV-1) isolate or a reptarenavirus mix, comprising UGV-1, University of Helsinki virus 1 (UHV-1) and Aurora borealis virus 1 (ABV-1) at approximately equimolar concentrations^31,49^ were used. The virus mix was prepared by inoculating I/1 Ki cells at a multiplicity of infection (MOI) of 0.1 to 0.01, and by pooling the supernatants collected at three, six, nine, and twelve days post-infection (dpi). The mix was stored in aliquots at −80°C, MOI of 1 to 10 was used in the infection experiments.

### Plasmids and molecular cloning

The open-reading frames (ORFs) for wild-type (wt) UGV-1 NP (Gene ID: 37629387), UHV-1 NP (GeneID:18821736) and HISV-1 NP (Gene ID: 41324517) were cloned into the pCAGGS-FLAG vector in frame with the C-terminal FLAG tag; the wtJUNV (Gene ID: 2545643) and wtLCMV (Gene ID: 956592) NP in pCAGGS-HA vector, in frame with the C-terminal HA tag^22^. wtJUNV NP, wtLCMV NP, and the pCAGGS-FLAG and pCAGGS-HA vectors were kindly provided by Prof. Luis Martinez-Sobrido (Texas Biomedical Research Institute, TX, USA) and have been described earlier^22,66^. The N-terminal mutations to the putative mitochondrial targeting signal (MTS) of UGV-1 NP (R6E, K15E, K16E, K17E, K20E) and JUNV NP (K5E, R11E, R17E, R18E) were generated using synthetic genes from Invitrogen which were subcloned into either pCAGGS-FLAG or pCAGGS-HA vectors in frame with the tag.

The NP ORFs of UGV-1 (wt and MTS-mutated), UHV-1, HISV-1, JUNV (wt), and LCMV were also subcloned into the pCR4Blunt-TOPO vector using the Zero Blunt TOPO PCR Cloning Kit for Sequencing, with One Shot TOP10 Chemically Competent *E. coli* (Thermo Fisher Scientific) following the manufacturer’s instructions. Individual clones were sent for Sanger sequencing at Microsynth Ag, and the plasmids containing the insert in coding orientation under the T7 RNA promoter were used for *in vitro* transcription/translation for the expression of radiolabelled NPs to assess mitochondrial import. The inserts were also subcloned into the pGEM4Z vector in coding orientation under the SP6 promoter (UGV-1, UHV-1, HISV-1, JUNV, and LCMV NP), or under the T7 promoter for HISV-1 NP. The molecular cloning followed standard procedures, and the primers used are listed in Supplementary Table 3.

### Transfections

The transfections were performed as described^67^. Lipofectamine 2000 (Invitrogen) was used for snake cells and FuGENE HD (Promega) for mammalian cells. Fresh conditioned medium was provided to the cells at 16-20 h post-transfection (hpt), and the analyses (immunoblot, immunofluorescence (IF) and transmission electron microscopy (TEM)) were performed on cells collected at three days post-transfection (dpt).

### Protein extraction from cultured cells

Trypsinized cells were washed twice with ice-cold PBS, pelleted by centrifugation (800 x *g*, 7 min, 4°C) and resuspended in ice-cold radioimmunoprecipitation assay (RIPA) buffer (50 mM Tris, 150 mM NaCl, 1% Triton X-100, 0.5% Sodium deoxycholate, 0.1% sodium dodecyl sulphate [SDS], pH 8.0) supplemented with complete protease inhibitor cocktail (Roche). Following 30 min incubation on ice and 10 min sonication, the lysates were clarified by centrifugation (13000 x *g*, 10 min, 4°C). The protein concentration was measured using Pierce BCA Protein Assay Kit (Thermo Fisher Scientific), and the lysates stored at −80°C.

### Subcellular fractionation and isolation of mitochondria from cultured cells

Subcellular fractionation was done using the Cell Fractionation kit-Standard (Abcam ab109719) following the manufacturer’s instructions. For the experiment, the cells were seeded on 10 cm diameter cell culture dishes (2.5*10^6^ cells/dish). The cells were inoculated with the reptarenavirus mix (final dilution 1:70, corresponding roughly to MOI of 5-10) immediately after the cells were seeded; non-inoculated cells served as controls. The cells for isolation were harvested at either one or two dpi. Before analyses, the cytosolic and mitochondrial fractions were 10-fold concentrated using Amicon Ultra-0.5 Centrifugal Filter Units (Merck Millipore) following the manufacturer’s instructions.

For isolation of mitochondria, confluent layers of cultured cells (surface area ≥150 cm^2^) were trypsinized, washed twice with ice-cold PBS, and pelleted by centrifugation (800 x *g*, 7 min, 4°C). The cell pellets were resuspended in 4 ml of ice-cold Mitochondria Isolation Buffer (MIB: 20 mM Hepes, 220 mM Mannitol, 70 mM sucrose, 1 mM ethylenediaminetetraacetic acid [EDTA], pH 7.6) with 2 mg/ml fatty acid-free bovine serum albumin (BSA, Sigma), supplemented with EDTA-free complete protease inhibitor cocktail (Roche) and kept on ice for 15-20 min. The cell suspensions were transferred to 5 ml Dounce homogenizers and manually homogenized. The homogenates were centrifuged (800 x *g*, 5 min, 4°C) and the pellets subjected to another round of homogenization and centrifugation. The supernatants were transferred into new tubes and the mitochondria pelleted by centrifugation (10,000 x *g*, 10 min, 4°C). The pelleted mitochondria were washed with 4 ml of ice-cold MIB, re-pelleted by centrifugation (10,000 x *g*, 10 min, 4°C), and re-suspended in 80-600 μl of MIB.

### Sodium dodecyl sulphate-polyacrylamide gel electrophoresis (SDS-PAGE) and immunoblot

The proteins from cell lysates and isolated mitochondria were analysed by separating 10-40 μg of protein (concentrations determined using Pierce BCA Protein Assay kit, Thermo Fisher Scientific), by SDS-PAGE. The samples were diluted in Laemmli Sample Buffer (LSB, final concentration: 0.3% SDS, 60 mM tris-HCl pH 6.8, 10% glycerol, 0.62% β-mercaptoethanol, 1% bromophenol blue) and denatured (10 min at 70°C), then loaded on 7.5% or 12% SDS-PAGE gels. After SDS-PAGE under standard conditions, the proteins were wet transferred (either 400 mA for 2 h or 160 mA overnight) onto nitrocellulose membrane (0.45 μm, Amersham Protran) in transfer buffer (25 mM tris, 200 mM glycine, 20% methanol). The membranes were blocked in Tris-buffered saline (TBS)-T (50 mM tris, 150 mM NaCl, 0.05% Tween 20, pH 7.4) with 5% (w/v) BSA. After primary (overnight at 4 °C or 2-3 h at room temperature (RT)) and secondary (2 h at RT) antibody incubations, and appropriate washes (three to five times for 5 min with TBS-T after antibody incubations, and twice with TBS before detection), the results were recorded using the Odyssey infrared imaging system (LI-COR Biosciences).

For the mitochondrial import assay, arenaviral NP samples were separated on pre-cast 7.5% SDS-PAGE gels (Bio-Rad). The mitochondrial import assay results were analysed by visualizing the proteins with the Coomassie stain (PhastGel Blue R, Sigma). The proteins were then fixed (20% ethanol and 10% acetic acid), treated with Amplify (Amersham) for 30 min and dried. Lysates obtained from import experiments performed on Su9-DHFR control were separated on 14% SDS-PAGE gels and transferred to nitrocellulose as described above. The ^35^S–methionine-labelled proteins on either dried gels or nitrocellulose membranes were visualized by autoradiographic detection using Amersham Hyperfilm MP (GE Healtcare).

### Antibodies

#### Primary antibodies

mouse anti-α-tubulin (Calbiochem CP06, dilution 1:500 in immunoblot), rabbit anti-mitochondrial outer membrane translocase 20 (TOM20, Santa Cruz sc-11415, dilution 1:1000 in immunoblot), rabbit anti-cytochrome c oxidase subunit IV (COX IV, Abcam ab16056, dilution 1:2000 in immunoblot), mouse anti-voltage-dependent anion-channel (VDAC, Abcam ab14734, dilution 1:500 in immunoblot), rabbit anti-mitochondrial ribosomal protein S35 (MRPS35, Proteintech 16457-1-AP, dilution 1:1000 in immunoblot), mouse anti-mitofusin 2 (MFN2, Abcam ab56889, dilution 1:200 in immunoblot), mouse anti-mitochondria “mtCO2” (Abcam ab3298, clone mtCO2, dilution 1:1000 in IF), mouse anti-FLAG (Flarebio CSB-MA000021M0m, dilution 1:500 in immunoblot and 1:1000 in IF), rabbit anti-FLAG (Flarebio CSB-PA000337, dilution 1:1000 in IF), mouse anti-HA (Sigma, H3663, dilution 1:500 in immunoblot and 1:1000 in IF), rabbit anti-HA (Flarebio, CSB-PA275079, dilution 1:1000 in IF), affinity-purified rabbit anti-UHV-NP^49^ (dilution 1:500 in immunoblot), rabbit anti-pan reptarenavirus NP^39^ (anti-pan-RAVs NP, dilution 1:6000 in IF and 1:1000 in immune EM). The polyclonal rabbit antiserum against the putative MTSs of arenavirus NPs (named MTS-NP) was raised against a synthetic multiepitope protein with the following sequence: MAALQRAAVNQLALKKKLNKMLSPFQRELNNQIFGGGGGMAALQEAAVNQLALEEELNEMLSPFQEELNNQIFGGGGGMSLSKEVKSFQWTQALRRELQGGGGGMSLSEEVESFQWTQALEEELQGGGGGMAAFQKAAVNQLALKKKLNKMLAPYQRELNNQIFGGGGGMAAFQEAAVNQLALEEELNEMLAPYQEELNNQIFGGGGGMAHSKEVPSFRWTQSLRRGLSGGGGGMAHSEEVPSFEWTQSLEEGLSHHHHHH. The polyclonal rabbit antiserum against hartmanivirus NPs was raised against a synthetic protein bearing amino acid stretches of HISV-1 NP with highest similarity to the NPs of other hartmaniviruses, sequence: EVLTNQLQVDYLFILIFCAKKQNMDLEALLELSGRCKLIFNKLPFTQKVLTQLSKSAKI ESSIEDLVIFTQTGYLDEKYLRKQGSGKLAGFMAKQHGMTKECKHAAKGGGGGGYSILREIENNLVLHDSPFRLNRQRFQSAVSALTGCVSDRMVSSGGGGGCKHKDGITVNTSEGSTTTYELLLHSILTTPTINAKIKNRTNVRRNGLNTVRFIGGGHHHHHH. Synthetic genes (in pET20b(+) vector) encoding the proteins were ordered from GeneUniversal, and the proteins produced in *E.coli* and purified as described^49,68^. Immunizations and sera collections were performed by BioGenes GmbH as described^49,68,69^. The resulting rabbit anti-MTS-NP was used at dilution 1:200 in both immunoblot and IF, and the rabbit anti-hartmanivirus NP (anti-Hartmani NP) antiserum at dilution 1:500 in immunoblot.

#### Secondary antibodies

IRDye 800CW Donkey anti-rabbit and IRDye 680RD Donkey anti-mouse (LI-COR Biosciences, both dilutions 1:10000 in immunoblot), Alexa Fluor 594 goat anti-rabbit (Thermo Fisher Scientific, dilution 1:400 in IF), Alexa Fluor 488 goat anti-mouse (Thermo Fisher Scientific, dilution 1:500 in IF), 18 nm gold-conjugated goat anti-rabbit IgG antibody (Milan Analytica AG, Rheinfelden, Switzerland; dilution 1:20 in immune EM).

### Submitochondrial localization assay

The submitochondrial localization assay used in the study is based on protease accessibility and was performed as described^48,70^, with some modifications. Freshly isolated mitochondria from a confluent cell layer (≥ 750 cm^2^) were resuspended in MIT buffer (320 mM sucrose, 10 mM tris, 1 mM EDTA, pH 7.4). The protein concentration was assessed using the Pierce BCA Protein Assay kit (Thermo Fisher Scientific). The mitochondria were pelleted by centrifugation (14,000 x *g*, 10 min, 4°C), resuspended in MIT buffer to yield 10 μg/μl, and divided into 4 different fractions of 100 μl each: A, B1 and B2, and C. Subsequently, pelleted mitochondria (14,000 x *g*, 10 min, 4°C) of each fraction were resuspended in 400 μl MIT for fraction A, 400 μl sonication solution (500 mM NaCl and 10 mM tris, pH 7.4) for fraction B1, 400 μl sonication solution and 3.8 μg/μl of proteinase K (proK, Recombinant PCR Grade [Roche]) for fraction B2, and 400 μl MIT-T (MIT + 0.5% Triton-X-100) for fraction C. The fractions B1 and B2 were sonicated on ice (three times for 30 s with 40% duty cycle in a Bandelin sonopuls sonicator) and fraction C was mixed by pipetting the solution up and down for 50 times. Fraction A represents intact mitochondria, fractions B1 and B2 vesicles obtained by sonicating mitochondria, and fraction C lysed mitochondria.

Fractions A and C were divided into four equal volume samples of 50-100 μl each. From fraction B1, a volume of 50-100 μl was taken and fraction B2 was divided into three equal volume samples of 50-100 μl each. MIT buffer (1/20 volume) without or with 30 μg/ml, 60 μg/ml or 120 μg/ml of proK was added to the samples, and after 20 min incubation on ice phenylmethylsulfonyl fluoride (PMSF, Sigma) was added to a final concentration of 2 mM, followed by 10 min incubation on ice. Then, 1% sodium deoxycholate was added to each sample to a final concentration of 0.05%, followed by 20 min incubation on ice. The proteins were precipitated by adding (1/6 volume) 100% (w/v) trichloroacetic acid (TCA), followed by 30 min incubation on ice and centrifugation (14,000 x *g*, 30 min, 4°C). The pelleted proteins were washed with 1 ml of ice-cold acetone, pelleted by centrifugation (14,000 x *g*, 30 min, 4°C), dried for 5 min at 37°C, and resuspended in 2x LSB to reach final concentration of 2.5 μg/μl. The assay was completed by analysing the samples via SDS-PAGE and immunoblot procedures, as described above.

### Immunofluorescence (IF) staining

For IF staining cells were seeded onto 13 mm coverslips (Thermo Fischer Scientific) in 24-well plates. The reptarenavirus-infected (MOI of 1-10) or reptarenavirus NP-transfected cells were PBS washed twice (at three dpi/dpt) and fixed with 4% PFA (in PBS) for 10-30 min at RT. After a PBS wash, cells were permeabilized and blocked (0.25% Triton-X 100 and 0.5% (w/v) BSA in PBS) for 5-10 min at RT, and washed twice with PBS. The cells were incubated overnight at 4°C with the primary antibodies, washed three to five times with PBS, incubated 1 h at RT with the secondary antibodies, washed three to five times with PBS, incubated 15 min at RT with DAPI (Novus Biologicals, 1 μg/μl, diluted 1:10000 in methanol), and washed twice with milli-Q water prior to mounting with FluoreGuard mounting medium (Scytek Laboratories). All primary and secondary antibodies were diluted in Dako REAL antibody diluent (Agilent technologies). Images were captured and analysed using a Nikon Eclipse TI microscope with NIS-Elements Microscope Imaging Software (Nikon).

### Transmission electron microscopy (TEM) and immune EM

TEM and immune EM studies were performed on cells grown in chamber slides (ibidi) as described^16^ and brain samples collected from a euthanized *Boa constrictor* with BIBD immediately after the animal’s death. Briefly, pelleted cells / tissue specimens were fixed in 1.5% / 2.5% glutaraldehyde, buffered in 0.2 M cacodylic acid buffer, pH 7.3, for 12 h at 5°C and routinely embedded in epoxy resin. Toluidin blue stained semithin sections (1.5 μm) and, subsequently, ultrathin (100 nm) sections were prepared and the latter contrasted with lead citrate and uranyl acetate and examined with a Philips CM10 transmission electron microscope at 80kV.

For immune EM, ultrathin sections were incubated for 30 min at RT in PBS with 1% BSA, followed by overnight incubation with primary antibody (diluted in PBS with 1% BSA) at 4°C. After washing with PBS, sections were incubated with the secondary antibody (diluted in PBS with 1% BSA) for 2 h at RT. Sections were then contrasted and examined as described^16^.

### Import of radiolabelled proteins into isolated mitochondria

The hybrid mitochondrial protein Su9-dihydrofolate reductase (DHFR), comprising the fusion of MTS of Subunit 9 of mitochondrial ATPase (Su9) of *N. crassa* with the DHFR of *M. musculus*, served as positive control for the protein import into mitochondria. Su9-DHFR was expressed under the SP6 RNA promoter in the pGEM4Z vector^50,51^, kindly donated by Dusanka Milenkovic (Max Planck Institute for Biology of Ageing, Cologne, Germany).

The mitochondrial protein import assay was carried out as described^52^, with some modifications. Full-length (tagged or untagged) NPs and Su9-DHFR were *in vitro* translated using rabbit reticulocyte lysate with either the TnT SP6 quick couple Transcription/Translation System (Promega) for cDNA sequences flanking the SP6 promoter, or the TnT T7 quick couple Transcription/Translation System (Promega) for cDNA sequences flanking the T7 promoter, following the manufacturer’s instructions, in the presence of 20 μCi [^35^S]methionine (Hartmann Analytic). The mitochondria were freshly isolated for the experiments from I/1 Ki, VI/1 Hz or Vero E6 cells. Protein concentrations were determined by the Pierce BCA Protein Assay kit (Thermo Fisher Scientific). The mitochondria were divided into 75 μg preparations, each one for a specific incubation time-point ± proK and/or CCCP addition, pelleted (10,000 x *g*, 10 min, 4°C), resuspendend in 100 μl import buffer (250 mM sucrose, 5 mM magnesium acetate, 80 mM potassium acetate, 20 mM Hepes-KOH, pH 7.4) supplemented with freshly added 10 mM sodium succinate, 5 mM adenosine triphosphate (ATP), and 1 mM dithiothreitol (DTT), and 5 μl of *in vitro* translation mix added. Carbonyl cyanide m-chlorophenyl hydrazone (CCCP) used at 1 mM or 2 mM concentration served as import blocker. The import reactions were performed at 30°C or 37°C under continuous rotation, and at the end of incubation proK was added (final concentration 50 μg/ml) to degrade the non-imported proteins. After 15 min incubation on ice, proK was inactivated by adding PMSF at a final concentration of 2 mM followed by 10 min incubation on ice. Next, the mitochondria were centrifuged (12,000 x *g*, 5 min, 4°C), washed with 200 μl SET buffer (250 mM sucrose, 10 mM tris-HCl pH 7.6, 1 mM EDTA, 0.1 mM PMSF), centrifuged (12,000 x *g*, 5 min, 4°C), and resuspended in 20 μl of 1x LSB. Samples were loaded on SDS-PAGE gels and analysed as described above.

### Bioinformatics

MitoProt II (http://ihg.gsf.de/ihg/mitoprot.html) and TargetP 1.1 (http://www.cbs.dtu.dk/services/TargetP-1.1) served for determining the probabilities for mitochondrial localization and the predictions for MTS cleavage sites for arenavirus NPs.

## ACKNOWLEDGEMENTS

This work was supported by the Academy of Finland (to J.H., grant numbers 308613 and 314119) and a grant from the Foundation for Research in Science and the Humanities at the University of Zurich to A.K.

The authors are grateful to the laboratory staff of the Institute of Veterinary Pathology, Vetsuisse Faculty, University of Zurich, for excellent technical support. We acknowledge also the laboratory support of the Institute of Veterinary Physiology and the Institute of Veterinary Pharmacology and Toxicology, Vetsuisse Faculty, University of Zurich. We would like to thank Dusanka Milenkovic (Max Planck Institute for Biology of Ageing, Cologne, Germany) for providing the Su9-DHFR construct in pGEM4Z vector. We thank Prof. Luis Martinez-Sobrido (Texas Biomedical Research Institute, TX, USA) for providing the expression plasmids for LCMV and JUNV wtNPs, and pCAGGS-HA and −FLAG vectors. We thank Prof. Nils-Göran Larsson (Karolinska Institute, Stockholm, Sweden), for providing us with some of the antibodies used to detect mitochondrial proteins. The laboratory work was partly performed using the logistics of the Center for Clinical Studies at the Vetsuisse Faculty of the University of Zurich.

## AUTHOR CONTRIBUTIONS

F.B., U.H., A.K. and J.H. conceived and designed the study. F.B. and U.H performed the experiments. F.B., U.H, A.K. and J.H. analysed the data. L.N. performed TEM and immune EM processing and acquisitions. F.B., U.H., A.K. and J.H. wrote the manuscript.

## SUPPLEMENTARY INFORMATION

Supplementary information accompanies this paper: Supplementary Figures 1-10 and Supplementary Tables 1-2 are provided in a PDF file and Supplementary Table 3 is provided in a separate Excel file.

For bioinformatic predictions for mitochondrial localization of arenaviral NPs belonging to the four different genera (*Reptarenavirus, Hartmanivirus, Antennavirus and Mammarenavirus*), the bioinformatic programs TargetP 1.1 (http://www.cbs.dtu.dk/services/TargetP-1.1) and MitoProt II (http://ihg.gsf.de/ihg/mitoprot.html) were used.

TargetP 1.1 reported data consist in: percentage of mitochondrial localization (% mt), predicted N-terminal mitochondrial targeting signal (MTS) length (amino acids, aa) and molecular weight (Mw, kDa) if available, and the reliability class (RC) with classes from 1=maximal reliability to 5=minimal reliability.

Mitoprot II reported data consist in: percentage of mitochondrial localization (% mt), position (aa) of the cleavage site and molecular weight of predicted MTS (MTS Mw) when available. Note that the two employed softwares often produce predictions with differing probabilities.

